# Massively parallel *in vivo* Perturb-seq reveals cell type-specific transcriptional networks in cortical development

**DOI:** 10.1101/2023.09.18.558077

**Authors:** Xinhe Zheng, Boli Wu, Yuejia Liu, Sean K. Simmons, Kwanho Kim, Grace S. Clarke, Abdullah Ashiq, Joshua Park, Zhilin Wang, Liqi Tong, Qizhao Wang, Xiangmin Xu, Joshua Z. Levin, Xin Jin

## Abstract

Systematic analysis of gene function across diverse cell types *in vivo* is hindered by two challenges: obtaining sufficient cells from live tissues and accurately identifying each cell’s perturbation in high-throughput single-cell assays. Leveraging AAV’s versatile cell type tropism and high labeling capacity, we expanded the resolution and scale of *in vivo* CRISPR screens: allowing phenotypic analysis at single-cell resolution across a multitude of cell types in the embryonic brain, adult brain, and peripheral nervous system. We undertook extensive tests of 86 AAV serotypes, combined with a transposon system, to substantially amplify labeling and accelerate *in vivo* gene delivery from weeks to days. Using this platform, we performed an *in utero* genetic screen as proof-of-principle and identified pleiotropic regulatory networks of *Foxg1* in cortical development, including Layer 6 corticothalamic neurons where it tightly controls distinct networks essential for cell fate specification. Notably, our platform can label >6% of cerebral cells, surpassing the current state-of-the-art efficacy at <0.1% (mediated by lentivirus), and achieve analysis of over 30,000 cells in one experiment, thus enabling massively parallel *in vivo* Perturb-seq. Compatible with various perturbation techniques (CRISPRa/i) and phenotypic measurements (single-cell or spatial multi-omics), our platform presents a flexible, modular approach to interrogate gene function across diverse cell types *in vivo*, connecting gene variants to their causal functions.

## Introduction

Multicellular organisms are comprised of a myriad of cell types, each endowed with unique molecular and functional profiles with distinct susceptibilities to diseases. Despite remarkable advancements over the past decade in identifying the genetic underpinnings of human diseases and disorders – yielding growing lists of high-confidence risk genes – the precise cellular contexts and tissue specificity in which these genes exert their effects remain largely elusive. For example, large-scale initiatives have made significant strides in mapping the genetic landscape of neurodevelopmental disorders, such as autism spectrum disorder and developmental delay (Fu et al., 2022; Kaplanis et al., 2020). Yet, given the diverse cell types that constitute the nervous system, our understanding of how these genetic variants confer vulnerability to particular cell types or brain regions, and thereby influence phenotypic outcomes, remains limited.

The rapidly evolving genomic technologies have enabled genetic studies across a wide array of cell types. Single-cell genomics has enhanced our ability to profile the cell type and cell state diversity across species. Meanwhile, CRISPR (clustered regularly interspaced short palindromic repeats) technology offers programmable perturbations to experimentally test gene function with scale (Schraivogel et al., 2023). *In vivo* Perturb-seq uses a single-cell RNA-seq (scRNA-seq) readout in pooled CRISPR screens to assay transcriptomic changes in a systematic way: sampling the effect of each perturbation in each cell type. It has provided a scalable platform to dissect genetic mechanism with high-content, high-resolution transcriptomic readout in developing brains (Dvoretskova et al., 2023; Jin et al., 2020). Through gene expression analysis, this approach revealed convergent molecular networks and specific neuronal and glial cell types impacted by risk genes within the context of a developing brain. While most *in vivo* CRISPR screens use guide RNA (gRNA) abundance as a proxy for cellular proliferation or depletion (Chen et al., 2015; Chow et al., 2017; Keys and Knouse, 2022; Tian et al., 2023; Tyson et al., 2021; Wang et al., 2018; Wertz et al., 2020), changes of gRNA abundance do not fully capture the molecular consequences and cellular phenotypes, especially changes in rare cell types that may play a critical role in the disease pathology (Bock et al., 2022; Townsley et al., 2020).

At present, conducting high-content phenotypic screens *in vivo* remains challenging due to at least two hurdles: (1) the need to scalably label, perturb, and isolate enough cells from various cell types *in vivo*, which is harder than most *in vitro* screens, and (2) by multiplexing the screen and mixing perturbation agents (gRNAs), we face the challenge to then robustly capture and deconvolute each cell’s perturbation identity in the sparse single-cell omics data. To compound these challenges, most Perturb-seq applications commonly rely on lentiviral vectors, which are known to have limited *in vivo* penetration, transduction, and thermostability (Higashikawa and Chang, 2001), hampers systemic screens in hard-to-reach tissues such as the adult central and peripheral nervous systems.

Adeno-associated viruses (AAVs) are single-stranded DNA viral vectors, highly effective for systemic gene delivery to many tissues with minimal immune response (Samulski and Muzyczka, 2014). Recent directed evolution strategies have resulted in a rapidly expanding toolbox for gene delivery in rodent models and humans (Chen et al., 2022; Kuzmin et al., 2021). Thus, AAV presents a promising strategy for delivering genetic perturbations to a wide range of cell types *in vivo*. However, the conventional AAV expression tends to be transient, at a relatively low level, with subsequently dilutions through cell divisions because the transgene remains episomal rather than integrated into genomic DNA (Lang et al., 2019). Together, these pose a challenge to accurately recover the perturbation identity of sparsely labeled cells in pooled assays like Perturb-seq. Most critically, the slow onset of AAV-mediated expression – often taking days or even weeks – is suboptimal for studying dynamic gene functions in fast-evolving cellular contexts such as neurodevelopment. For example, peak AAV-transgene expression commonly occurs after more than seven days, a timespan that encompasses the entire window for corticogenesis. These limitations underscore the pressing need for an AAV system that delivers rapid and robust transgene expression, to expand the capabilities of Perturb-seq *in vivo*.

Here, we report our development of an AAV-based, massively parallel *in vivo* Perturb-seq platform to target a broad spectrum of tissues and cell types with gene expression-based characterization at single-cell resolution. Through a barcoded screen of 86 phylogenetically diverse AAVs *in vitro* and *in vivo*, we identified several serotypes including AAV-SCH9, which enable rapid and robust transgene delivery in newborn neurons and progenitors within 48 hours (versus 2-3 weeks by the state-of-the-art methods). We further combined this vector with a transposon system to ensure sustained expression of gRNAs in both target cells and their daughter cells, allowing efficient gRNA capture in the single-cell analysis. In our proof-of-principle *in utero* perturbation screen, we identified the cell type-specific impact of perturbations on diverse neuronal populations. Gene expression analysis elucidated cell type-differential transcriptomic changes: *Foxg1* predominantly affected Layer 6 corticothalamic (L6 CT) neurons, and its loss-of-function, strongly associated with neurodevelopmental delay, altered transcriptional networks, and led to hybrid cell fates. These effects are highly specific to Layer 6 CT neurons but not observed in other cell types from the same developmental lineage and layer. Our system can profile over *30,000* cells in one experiment – providing a massively parallel approach that is required for systematically investigating *hundreds* of risk genes in heterogeneous tissues (*hundreds* of brain cell types) with robust power. This is a modular and versatile strategy to efficiently study diverse cell types *in vivo*, paving the way to comprehensively approach the mechanism of panels of human disease risk genes in physiological contexts.

## Results

### Barcoded AAV screens identify serotypes efficiently expressed in the developing brain

Existing Perturb-seq systems rely on lentiviral vectors, which have relatively low packaging yield and limited tissue penetration *in vivo*. AAV vectors are advantageous for *in vivo* studies due to their high-titer production yields and capsid engineering potential. However, most AAVs take weeks to reach peak expression, which is suboptimal for transducing newborn neurons and progenitors with high expression level in a rapidly developing brain. To identify an AAV serotype targeting the developing brain *in vivo*, we constructed a barcoded library of 86 phylogenetically diverse AAV serotypes; each expressing a green fluorescent protein (GFP) with a unique DNA barcode upstream of the polyadenylation sites (**Fig. 1A**). This library consists of engineered serotypes that were reported or published previously, including AAV1, AAV2, AAV3B, AAV4, AAV5, AAV6.2, AAV7, AAV8, AAV9, AAV10, AAV 12 and AAV13; several of these serotypes are commonly used for *in vivo* studies with minimal toxicity as previously reported (Table S1).

To identify a fast-acting serotype in developing mouse brains (C57BL/6J strain), we administered this library of AAV vectors *in utero* into the lateral ventricles at embryonic day 13.5 (E13.5) (0.5-1.5μL, 0.5-1.5e^10^ viral genomes/embryo). At 48 hours, we observed many GFP^+^ cells in the cortex and ganglionic eminence (**Fig. 1B-C**). Through immunofluorescence marker labeling, we observed that 56% of GFP^+^ cells were newborn neurons (TBR1^+^) that our AAV library directly or indirectly transduced. We found 3% of GFP^+^ cells expressed the intermediate progenitor (IP) marker TBR2, which indicates that either the AAVs were less potent at transducing IPs, or the GFP was quickly diluted as IPs differentiated. At 24 hours, a minimal GFP signal was detected from the brain tissue with immunofluorescent amplification, suggesting that most AAV vectors took 24 to 48 hours to express detectable levels of the transgene *in vivo*. In parallel to this *in viv*o analysis, we conducted the screen *in vitro* with a mouse hippocampal neuronal cell line HT22 (Fig. S1A) and could detect GFP expression *in vitro* at 24 and 48 hours post transduction (Fig. S1B).

### AAV-SCH9 rapidly transduces developing brains within 48 hours

To identify the AAV serotypes that were enriched in GFP^+^ cells, we purified the cells from the neocortex (*in vivo*) and HT22 cells (*in vitro*) at 24-and 48-hours post-transduction and quantified the barcode abundance using next-generation sequencing (**Fig. 1D, S1C-H**). Compared to the initial distribution in the AAV library, 48 hours post-transduction, both *in vivo* and *in vitro*, we detected significant shifts in the barcode distribution. We found that, 48 hours post-transduction, several AAV serotypes were enriched *in vitro* (AAV-P1529, AAV2-NN, AAV2-P1583 and AAV-DJ), several serotypes were enriched *in vivo* (AAV1, AAV2-P1558, AAV-Hu48.2 and AAV2-7M8), and several were enriched in both (AAV-SCH9, AAV-SCH9repeat136bp, AAV2-P1576, AAV2-P1579 and AAV2-P1596) (**Fig. 1D, S1D-I, Table S1**). This likely reflects a shared pattern of AAV tropism for newborn neurons and progenitors *in vivo* and *in vitro*, as well as a distinct pattern of enrichment *in vivo*.

To further examine the pattern of AAV barcode distribution across time points and across different contexts, we performed a principal component analysis of the samples (**Fig. 1E**). PC1 explained 63% variance of the data and separated *in vitro* data (24 and 48h), *in vivo* data (48hr), from *in vivo* data (24h) and the initial AAV library. PC2 (10% variance) separated *in vivo* (48h) data from all other samples. Consistent with the *in vivo* tissue analysis, it takes 48h to observe the change of barcode distribution *in vivo*, while the impact is quicker and more consistent at 24h and 48h *in vitro*.

**Figure 1.**
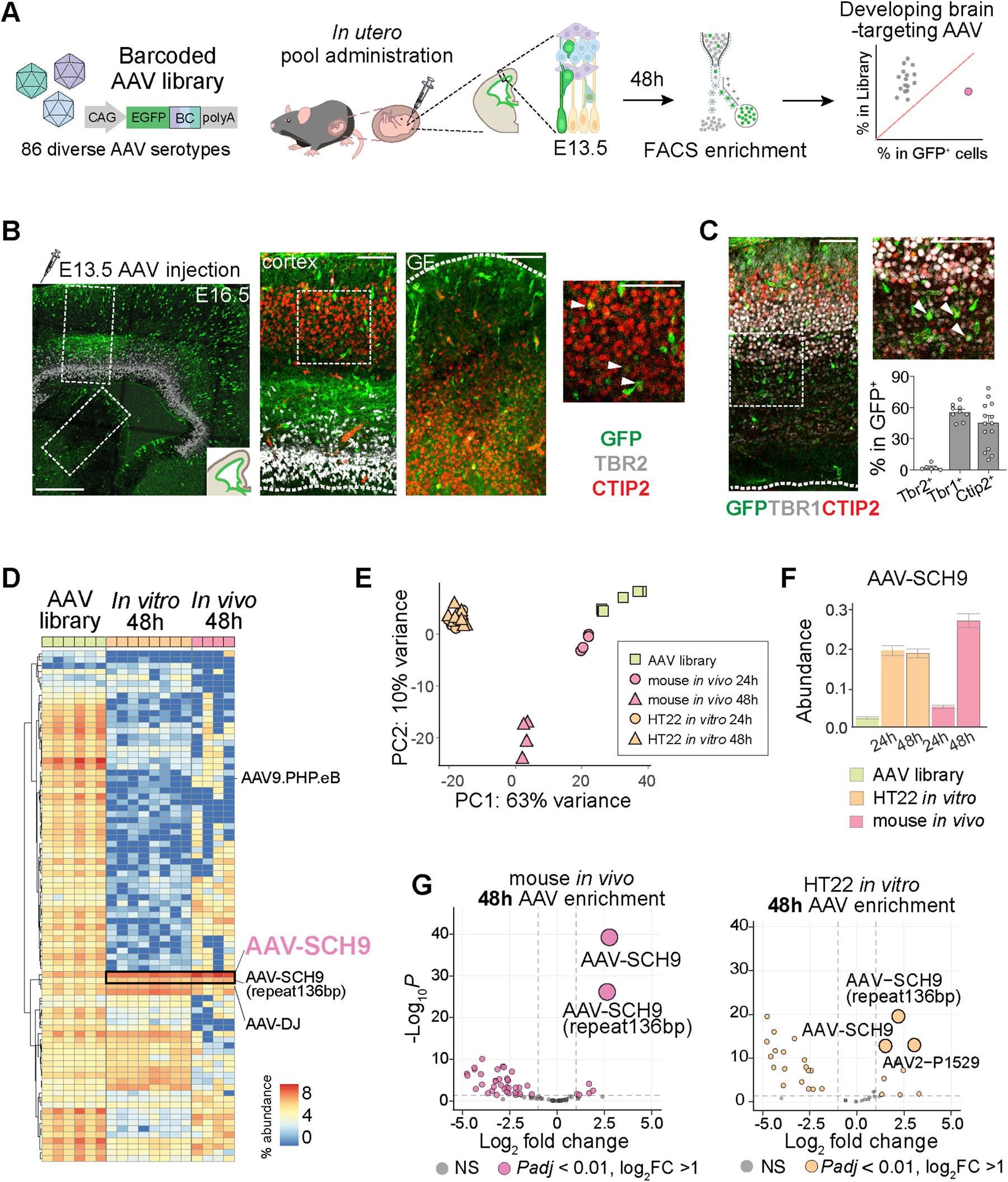
Barcoded AAV serotype screen *in vivo* identified AAV-SCH9 efficiently targeting developing brains. (A) Schematics of AAV library administration *in utero* at E13.5 followed by Fluorescence-activated cell sorting (FACS) cell enrichment and barcode analysis with next generation sequencing. (B-C) Immunofluorescence analysis of brain sections two days after AAV library administration: (B) co-stained with markers of newborn projection neurons (TBR1 and CTIP2) and intermediate progenitors (TBR2), in dorsal cortical laminar and ganglionic eminence (GE); (C) quantification of percentage of GFP^+^ cells co-expressing neuronal and progenitor markers including TBR1, TBR2 and CTIP2. (D) Heatmap of proportion of 86 AAV serotype abundance in AAV library, 48 hours post transduction in HT22 cells and in embryonic mouse brain; each row represents the abundance of an AAV serotype. (E) Principal component analysis of the percentage abundance of AAV library, HT22 cells and mouse brain 24-or 48-hours post transduction. (F) AAV-SCH9 percentage abundance in the initial AAV library (input for transductions), HT22 cells and mouse brain 24-or 48-hours post transduction. Error bars indicated standard error of the mean. (G) Volcano plots of AAV serotype changes in mouse brain (left) or HT22 cells (right) 48 hours post transduction compared to the initial AAV library. Scale bars indicate 250μm (left in B), 50μm (right in B and in C).

We observed a 11.5-fold increase in the proportion of the barcode pool in our top hit, AAV-SCH9: it constituted 2.3% of the initial AAV library and 27.4% of the enriched population 48 hours after *in vivo* transduction (**Fig. S1D**). Differential expression analysis revealed that AAV-SCH9 was the most significant hit from the *in vivo* experiment, followed by a variant of the same serotype (AAV-SCH9-repeat136bp). Both AAV-SCH9 and its variant showed significant enrichment *in vitro* 24 hours post transduction that persisted at 48 hours (**Fig. 1G, S1D-F**). We also quantified the relative expression dynamics of several top hits, including AAV-SCH9, by comparing the abundance of the barcodes in 24 and 48-hour post-transduction (**Fig. 1F, Fig. S1I**).

Critically, compared to AAV-SCH9, several widely used neuron-targeting AAVs required longer to reach peak expression. Within the initial 48 hours, AAV9-PHP.eB and AAV-DJ showed only minimal expression *in vivo*, with 0.2-fold and 1.1-fold changes relative to their compositions in the initial AAV library, respectively (**Table S1, Fig. 1D**). By contrast, AAV-SCH9 demonstrated a 11.5-fold increase in expression abundance, making it an optimal choice for perturbing and studying gene function in a dynamic developmental context when cells are actively differentiating and maturing.

### Diverse cell type tropisms of AAV serotypes *in vivo* with single-cell resolution

We identified several serotypes, including AAV-SCH9, that exhibited efficient transduction of the developing brain and neuronal cell lines using bulk measurements, but their precise cell type-specificity was still unclear. To further characterize the tropism of the AAV hits, we selected the top 14 serotype hits to create a new, secondary library for validation with single cell resolution (**Table S1**). For these experiments, each AAV was designed to express a GFP reporter with one of a set of three unique barcodes upstream of the polyadenylation sites and then pooled with equal titer (**Fig. 2A, S2A**). We administered this 14-AAV library into the lateral ventricle of E13.5 mouse embryonic brain and collected cortical cells at E15.5 to perform droplet-based scRNA-seq with both the sorted (GFP^+^) and unsorted populations (**Fig. 2A-B**). We assigned the AAV serotype identities by using the barcodes, captured in a dial-out PCR library (**Fig. 2C**) (see Methods).

**Figure 2.**
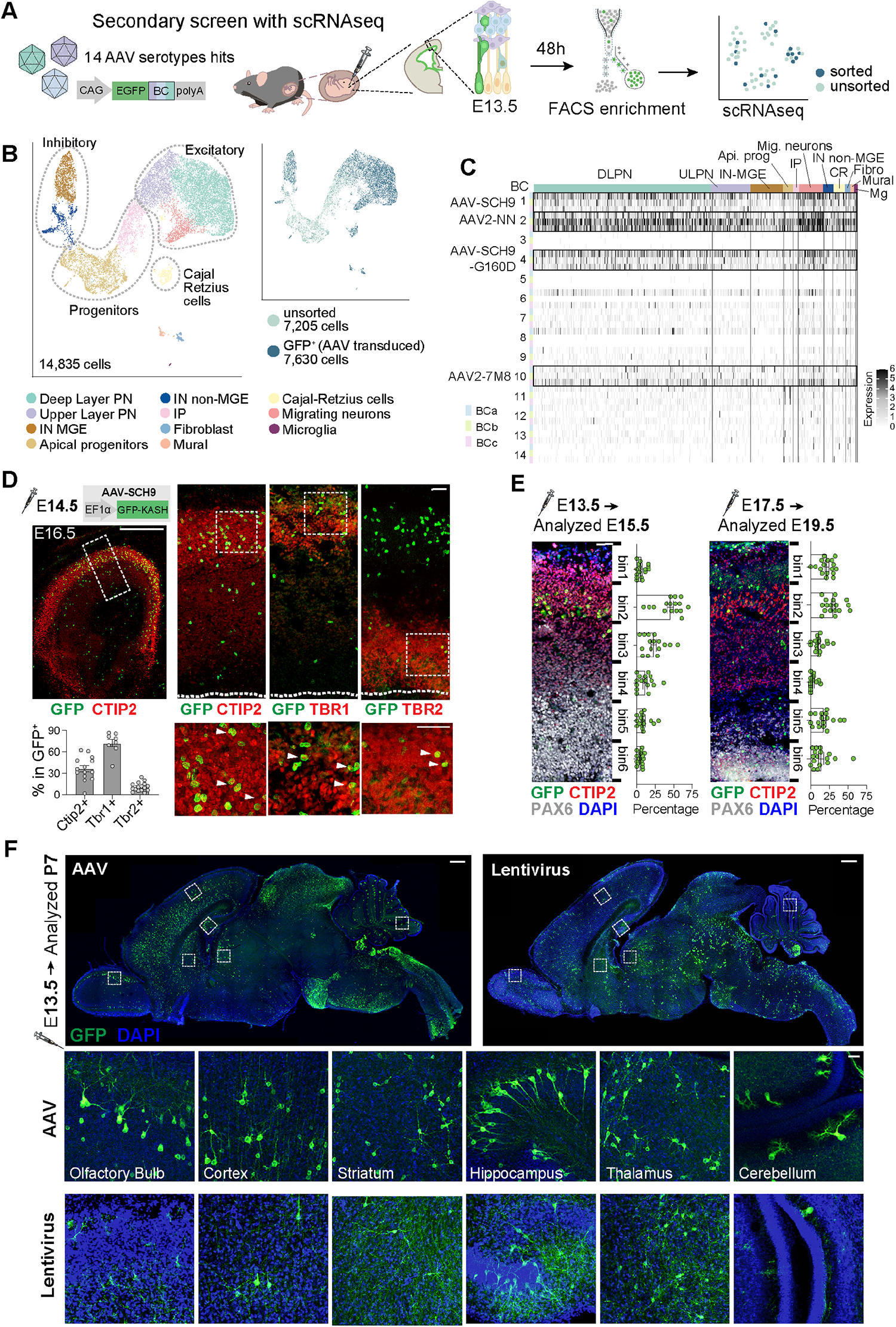
AAV-SCH9 labeled newborn neurons spreading across brain regions *in vivo* across developmental stages. (A) Schematics of a secondary AAV serotype screen: 14 AAV serotypes were barcoded and introduced in pool *in utero* at E13.5, followed by scRNA-seq 48 hours later. (B) Uniform Manifold Approximation and Projection (UMAP) visualization of 11 major cell populations identified (left) from sorted (GFP^+^) and unsorted cells (right); cell types include: upper and deep layer projection neurons (ULPN, DLPN), migrating neurons (Mig. neurons), apical progenitors (Api. prog), intermediate progenitors (IP), interneurons derived from the medial ganglionic eminence (IN-MGE), interneurons derived from the non-medial ganglionic eminence (IN-non-MGE), Cajal-Retzius cells (CR), fibroblast (Fibro), mural cells (Mural), and microglia (Mg). (C) AAV serotype barcode expression in each cell type. Each row represents an AAV barcode, and each AAV serotype is associated with a distinct set of 3 barcodes (BCa, BCb and BCc). Each column represents a given cell, with coloring based on the number of barcode UMI, arranged by cell types. (D) Immunofluorescence analysis of AAV-SCH9-GFP-KASH E14.5 transduced brain sections co-stained with markers including CTIP2, TBR1, and TBR2 as well as the percentage of marker co-expression in the GFP^+^ cells. Boxes on top panels indicate chosen fields of view in the bottom panels; arrows indicate representative cells with marker co-localizations. (E) AAV-SCH9-GFP-KASH was administered at two different time points (E13.5 or E17.5) and the cortical sections were analyzed 48 hours later to quantify GFP^+^ cells distribution across laminar layers, which were divided evenly into six bins from pia (bin 1) to ventricle (bin 6) (n=2-3 animals/condition). Each dot represents a crop of cortical column from a brain section. (F) AAV-SCH9-GFP-KASH or lentiviral reporter (GFP) administered at E13.5 resulted in diverse brain region labeling at P7. Scale bars indicate 500μm (left in D and top in F) or 50μm (right in D, E, and bottom in F).

After quality control (see Methods), we retained a total of 14,835 neocortical cells for further analysis (7,630 cells from the FACS-enriched GFP^+^ population, and 7,205 cells without enrichment) (**Fig. 2B, S2B-C)**. We partitioned the cells into major cell types and annotated them based on known marker gene expression (**Fig. S2E-F, Table S4**) (Di Bella et al., 2021; La Manno et al., 2021; Tasic et al., 2018). These cells were clustered into 11 cell types including upper and deep layer projection neurons (ULPN, DLPN), migrating neurons (Mig. neurons), apical progenitors (Api. prog), intermediate progenitors (IP), interneurons derived from the medial ganglionic eminence (IN-MGE), interneurons derived from the non-medial ganglionic eminence (IN-non-MGE), Cajal-Retzius cells (CR), fibroblast (Fibro), mural cells (Mural), and microglia (Mg) (**Fig. S2D**).

The relative abundance of most of the cell types in GFP^+^ and unsorted populations was similar. Two populations, apical and intermediate progenitors, showed decreased representation in the GFP^+^ population (**Fig S2G**), reflecting a reduced capacity of the AAV to transduce dividing cells, or cell division diluting the transgene products, or both. From inspecting the individual barcodes across cell populations, we found that four serotypes were highly efficient in transducing *in vivo*: AAV-SCH9 (BC1), AAV2-NN (BC2), AAV-SCH9-G160D (BC4) and AAV2-7M8 (BC10) (**Fig. 2C, S2H**). Within the AAV-SCH9 transduced populations, we detected 12.0% upper layer projection neurons, 56.5% deep layer projection neurons, 10.1% MGE-derived interneurons, 7.9% migrating neurons, 2.8% apical progenitors, and 1.3% intermediate progenitors (**Table S3-4**).

### *In situ* characterization of AAV-SCH9 reveal its fast-acting dynamics and broad neuronal labeling in the developing brain

Using bulk sequencing and scRNA-seq, we identified and validated AAV-SCH9 as an effective vector to rapidly (< 48 hours) transduce embryonic cortical tissues *in vivo*. Previously, AAV-SCH9 has been reported to target subventricular neural stem cells in the adult brain (Ojala et al., 2018). To further characterize its brain-wide action and tropism in development, we administered AAV-SCH9 expressing a nuclear membrane-anchored fluorophore (GFP-KASH) *in utero* into the lateral ventricles at embryonic day 14.5 (E14.5). After 48 hours, we observed many GFP^+^ cells in the neocortex, ganglionic eminence, and around the lateral ventricle, similar to the pattern from the 86-AAV library (**Fig. 2D and 1B-C**).

We co-stained the section for markers of deep layer excitatory projection neurons (CTIP2 and TBR1) and intermediate progenitors (TBR2). We found on average 37% of GFP^+^ cells express CTIP2 and 73% express TBR1, indicating that AAV-SCH9 induced broad expression in newborn neurons. For GFP^+^ cells, 11% expressed the intermediate progenitor marker TBR2, indicating either the transduction was less efficient in progenitors and/or the expression was quickly diluted as IPs divided. The relative proportion of cell types measured by immunohistochemistry and scRNA-seq data generally agree, as expected (**Fig. S2F, H, Table S5**). These data confirmed the ability of AAV-SCH9 to label neurons and intermediate progenitors *in vivo* within 48 hours.

AAV-SCH9 may have directly transduced neurons, or first transduced progenitors which then differentiated into neurons. To identify its tropism and differentiate these two possibilities, we performed *in utero* transduction of AAV-SCH9-GFP-KASH at two embryonic ages when two distinct populations of neurons (deep versus upper layers) are born. If the AAV only labels progenitors, we expected to observe enrichment of GFP expression in different layers, targeting different classes of newly born neurons on different injection days (only deep layers for E13 injection, and only upper layers for E17 injection, respectively). Brains were harvested after 48 hours and co-stained with DLPN marker CTIP2 and neural progenitor marker PAX6 to distinguish the cortical layers. We divided the cortical layers evenly into 6 bins from pia (bin 1) to ventricle (bin 6): the E13.5 transduced cells were enriched in bin 2 (46%) and bin 3 (23%) with CTIP2 expression, indicating their deep layer cortical sub-cerebral projection neuron identities (Arlotta et al., 2005; Chen et al., 2008). By contrast, the E17.5-transduced cells were broadly distributed in the later-born ULPNs in bin 1 (22%) and Layer 5 DLPNs in bin 2 (31%), indicating that AAV-SCH9 labeled newborn neurons. (**Fig. 2E**). Interestingly, we observed a higher level of GFP^+^ cell labeling in the ventricular and subventricular zone in the E17.5-than E13.5-transduced sample (18% versus 9% in bin 5, 15% versus 5% in bin 6), supporting the progenitor-labeling capacity of AAV-SCH9 and consistent with the slower cycling and differentiation capacity in the late neurogenesis stage (Greig et al., 2013). These data, together with the scRNA-seq results, collectively support that AAV-SCH9 transduced both newborn neurons and progenitors in embryonic mouse brain with a fast onset of expression.

One of the advantages of AAVs is their excellent tissue penetration and labeling *in vivo*. Besides the neocortex, AAV-SCH9 can transduce additional regions, including olfactory bulb, striatum, hippocampus, thalamus, and cerebellum, with labeling density suitable for future cross-region Perturb-seq studies (**Fig. 2F**). By contrast, we observed limited expression with lentiviral transduction (using optimal conditions, >9×10^9^ U/mL high-titer vector): there were much fewer GFP^+^ cells in most regions, especially cerebellum, thalamus, and striatum (**Fig. 2F**), possibly due to limited tissue penetration *in vivo*. Altogether, this scalable Perturb-seq platform holds greater potential for its ability to access abundant cells across brain regions, allowing a thorough comparison of the brain region-and cell type-specific perturbation effects of disease risk genes *in vivo*. Furthermore, AAV-SCH9 transduction surpasses conventional lentiviral vectors in labeling neocortex and other brain regions, significantly streamlining the sampling process for scRNA-seq and reducing the associated costs and labor required.

### Transposase hypPB stabilizes and enhances transgene expression

In contrast to lentiviral vectors, most AAVs remain episomal in host cells due to the lack of integrase (Deyle and Russell, 2009). Therefore, when performing Perturb-seq using AAV vectors in the rapidly dividing cellular context *in vivo*, the gRNA and fluorophore expression from AAV will be diluted, and eventually lost, upon cell division and growth. To overcome the challenge of gRNA dilution and improve genetic perturbation and labeling efficiency *in vivo*, we designed a dual-vector system including a hypPB (hyperactive piggyBac) transposase and a transposon with inverted repeats flanking the gRNA and fluorophore (Moudgil et al., 2020) (**Fig. 3A**). Upon transduction, hypPB could integrate the transgene into the nuclear genome for inheritance in the daughter cells, allowing consistent expression rather than transient, episomal expression (**Fig. 3A).**

We first tested the design *in vitro* and transfected HT22 cells followed by time-lapse imaging. In the presence of hypPB, we detected more GFP^+^ cells (**Fig. 3B**, S3A) and enhanced GFP expression levels with faster onset (Fig. S3B); the expression was stable with cell passages over the course of six days (Fig. S3C). hypPB allowed faster expression onset within a few hours post-transfection which would be highly advantageous for labeling and perturbing cells during the dynamic developmental process *in vivo*.

### hypPB improves transgene expression *in vivo* through embryonic transduction

To evaluate whether hypPB can enhance gene expression *in vivo*, we first administered AAV-SCH9 containing a transposon with a fluorophore, with or without an AAV-SCH9-hypPB *in utero* at E14.5 and performed immunofluorescence analysis at P7. We detected hypPB (HA-tagged) expression in many brain regions including cortical laminar layers, similar to the expected AAV-SCH9 tropism (**Fig. 3C**). hypPB expression did not introduce overt toxicity in development (**Fig. 3C, S3D**) based on the absence of obvious changes in the expression of gliosis markers GFAP and IBA1 in the presence of hypPB.

**Figure 3.**
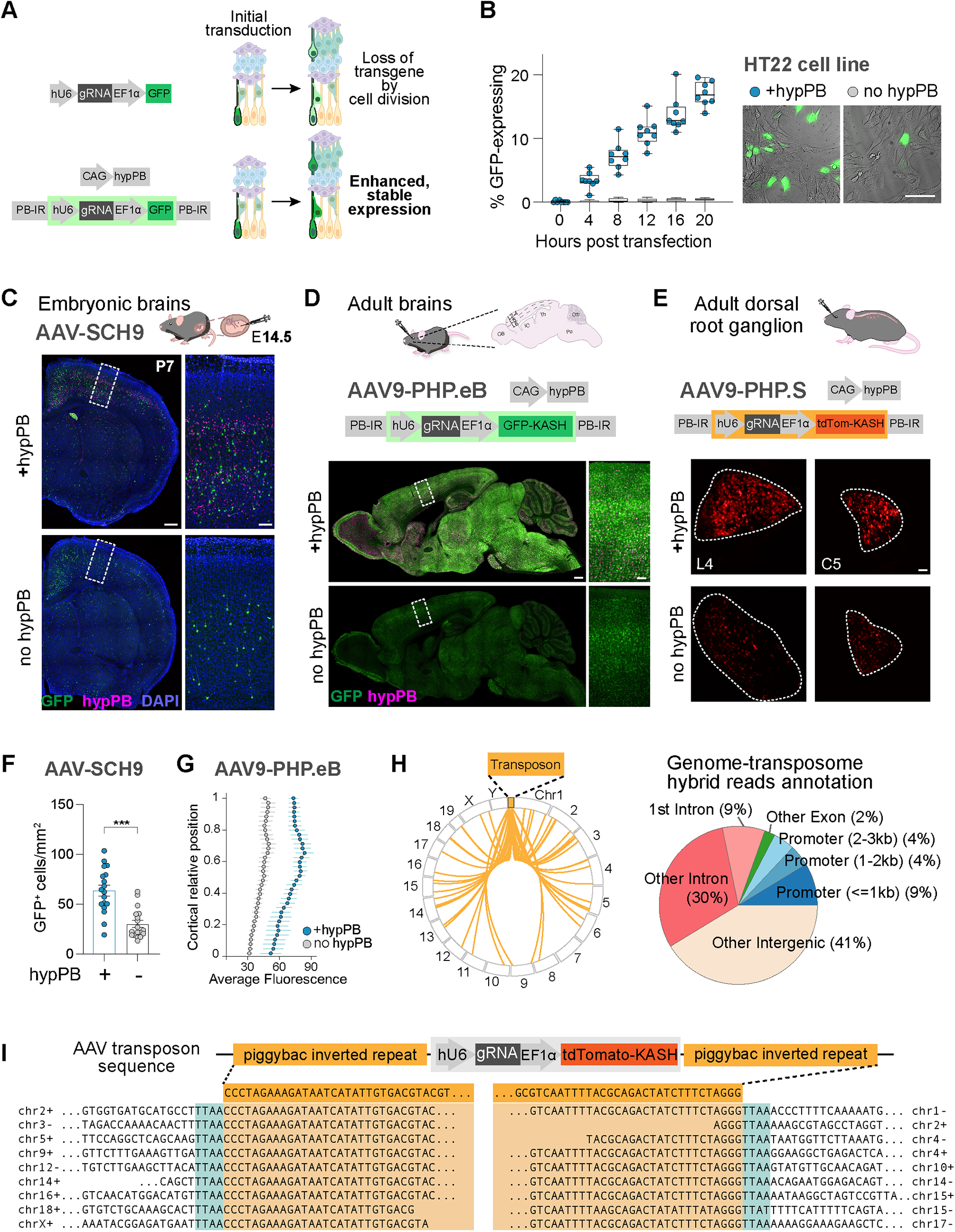
hypPB transposon enhanced and stabilized expression in embryonic and adult brains and peripheral nervous systems. (A) Schematics of the molecular design to enhance transgene expression and prevent loss due to cell division and differentiation. Shaded box indicates transposon flanked by the inverted repeats (IR). (B) Timelapse imaging showed co-transfection of hypPB increased the expression of the GFP transgene *in vitro*. Each dot represents analysis from a chosen field of view from a well, the Y axis represents the percent of cells expressing GFP. (C-E) Transposon stabilized expression *in vivo* across embryonic brain, adult brain, and adult dorsal root ganglion using three targeting vectors: AAV-SCH9, AAV9-PHP.eB and AAV9-PHP.S. (F) AAV-SCH9 labeled more cortical neurons in the presence of hypPB; Y axis indicated the number of GFP-expressing cells per mm^2^ (n=3 animals/condition). Asterisks indicate *P-value*<0.0001 with unpaired t-test. Each point represents a crop of cortical column from a brain section. (G) AAV9-PHP.eB labeled cortical neurons with increased expression intensity with hypPB: average fluorescence intensity (X axis) across cortical layers (Y axis), from ventricular zone (0) to pia (1) (n=3 animals/condition). (H) Whole genome sequencing of AAV-SCH9 transduced cells *in vivo* showed the genomic regions of integration events. Left: each line indicates a unique hybrid read between mouse genome and transposon, illustrated as a wheel plot. Right: plot of percentage of integration events that occurred in different intergenic and coding regions in the genome. (I) Example reads that include the junction of mouse genome and transposon, indicating the integration sites. Top: schematic illustration of the transposon sequence. Bottom: example reads aligned to the transposon sequence (in orange boxes) and to the mouse genome with their chromosome numbers. Reads on the left and right on the same line are unrelated to each other. Scale bars indicate 100μm (in B, right in C, right in D and in E) or 500μm (left in C and left in D).

We quantified the number of GFP^+^ cells in the somatosensory neocortex from brains administered with AAV-transposon, with and without hypPB, *in utero*. Co-transduction of hypPB increased the labeling efficiency: 2.2-fold more cells were GFP^+^ (**Fig. 3F**). Consistently, we divided the cortical laminar layers evenly into 4 bins and detected 3.0-fold more GFP^+^ cells with hypPB co-expression in the upper layers (15 versus 5 cells/mm^2^, corresponding to bin 1), 1.8-fold in Layer 5 (108 versus 59 cells/mm^2^ in bin 3), and 3.2-fold in Layer 6 (123 versus 39 cells/mm^2^ in bin 4) (**Fig. S3E**). This result is consistent with our *in vitro* study that hypPB co-expression led to retention of expression in dividing and differentiating cell lineages.

### hypPB enhances transgene expression in adult central and peripheral nervous system

The utility of applying Perturb-seq *in vivo* have been demonstrated in the past in embryonic brains (Dvoretskova *et al*., 2023; Jin *et al*., 2020). However, the need for a comprehensive *in vivo* screen platform for the adult central and peripheral nervous systems – especially pertinent to neurodegenerative diseases – still exists. We hypothesized that hypPB could amplify AAV transgene expression in postmitotic neurons in adult central and peripheral nervous systems, where AAV transgene levels have been observed to be relatively low compared to genomic expression (Lang *et al*., 2019). To test this hypothesis, we delivered AAV vectors to postmitotic neurons in adult mice using two variants that cross the blood-brain barriers: AAV9-PHP.eB for neurons and glia in the brain, and AAV9-PHP.S for peripheral neurons, both through retro-orbital administrations (Chan et al., 2017).

In adult brain, hypPB generally increased GFP expression levels (**Fig. 3D**). In the somatosensory cortex, average fluorescence intensity was 1.7-fold higher across laminar layers with hypPB (**Fig. 3G**). Furthermore, by extracting nuclei from the cortex and performing flow cytometry analysis, we found that the AAV9-PHP.eB transposon system can label 6.5-7.0% of total nuclei in the cortex, 4.0-fold higher than without hypPB **(Fig. S3F)**. This indicates hypPB can enhance transgene expression in adult, post-mitotic cells. Notably, different from the AAV-SCH9 delivery, elevated levels of gliosis markers IBA1 and GFAP were detected in the retro-orbital AAV9-PHP.eB-hypPB transduction conditions **(Fig. S3G)**. This is likely due to the neuroinflammation response by the systemic viral administration method and the amount of vector used (Perez et al., 2020) as well as hypPB genome integration, indicating the importance of carefully evaluating AAV administration to minimize toxicity concerns.

To test the performance of this system in the adult peripheral nervous system including the dorsal root ganglion, we performed retro-orbital injection of the AAV9-PHP.S gRNA-tdTomato construct, with or without the AAV-hypPB (**Fig. 3E**). Similarly, the presence of hypPB also increased the reporter expression levels and labeling efficiency (**Fig. 3E**). Altogether, hypPB transposon can further enhance transgene expression in adult nervous systems, combined with other AAV9 variants. This opens doors to study genetic perturbations and gene function in adult tissues, especially in the context of aging or degeneration of central and peripheral nervous systems, beyond the capability of conventional lentivirus-based genetic screens.

### hypPB transposon integrates into the host genome mostly in non-exonic regions

Genome integration of the transgenes, via lentiviral vectors or transposons, could trigger unintended changes in the genome and cellular activities. For instance, integrations close to coding regions could profoundly influence the function of the affected gene. To characterize the transposon integration preferences in neurons *in vivo*, we extracted the mouse genomic DNA from over 170,000 tdTomato^+^ nuclei from brains co-administered with AAV-SCH9-hypPB and transposons *in utero*. We then performed 60x whole-genome sequencing to identify hybrid reads that captured the junctions between the mouse genome and transposon, which provides evidence of transposon integration sites (**Fig. 3H-I**) (see Methods). We identified 46 such insertion events, several of which were supported by multiple reads, distributed across the mouse genome (**Table S6**). Overall, most of the insertion events were in the intergenic regions (41%). We observed a slight preference for transcription start sites amongst insertions within the promoter regions (**Fig. 3H-I**), consistent with the literature on hypPB specificity (Chen et al., 2020). Integration events occurred at the expected TTAA flanking sites, further demonstrating the expected insertion performance of hypPB *in vivo*. These analyses showed that hypPB integration sites are widely dispersed across the genome. The perturbation effects in each cell could be confounded by the issue of hypPB-induced genome integration, which also exists for lentiviral vector-based screens. Since the events are largely random and in the intergenic regions, sampling large numbers of cells for each perturbation group could minimize this bias to enable a robust and well-powered analysis.

### AAV-hypPB permits gRNA capture with sparse scRNA-seq readout

One challenge of *in vivo* CRISPR screens is to retain high gRNA expression for efficient gene editing as well as gRNA recovery in the sparse scRNA-seq data (Kalamakis and Platt, 2023). We next tested if the transposase system could also enhance gRNA expression levels and found that *in vitro* hypPB co-transfection led to a 4.2-fold increase in the gRNA expression detected **(Fig. S4A)**. Interestingly, the presence of Cas9 stabilized gRNA expression by 5.7-fold, likely by forming ribonucleoprotein complexes to prevent gRNA degradation (Hendel et al., 2015). With the co-expression of hypPB and Cas9, gRNA expression level was increased by 10.8-fold, likely through additive mechanisms, which could improve the gene editing efficiency as well as gRNA detection in the scRNA-seq readout.

With the fast-acting AAV-SCH9 and the hypPB system to enhance the expression, we moved on to characterize the gRNA identity recovery rate of this Perturb-seq platform by two established scRNA-seq strategies. Embryonically transduced cortical cells were postnatally dissociated, purified and followed by scRNA-seq, with two methods to capture gene expression and gRNA identities: 3’ scRNA-seq, which uses on-beads barcoded oligo-dT and gRNA primers to label and amplify the 3’ ends, or 5’ scRNA-seq, which uses in-solution oligo-dT and gRNA primers to capture the transcript from the 5’ end (Replogle et al., 2020). The 5’ method benefits from a higher concentration of gRNA-specific primers in solution (rather than on the beads) during reverse transcription, which might be expected to give rise to a higher gRNA capture rate.

We observed that, with similar sequencing depths, the 5’ and 3’ scRNA-seq gene expression analysis resulted in similar cell clustering and data quality (**Fig. S4C-D, Table S7**). However, with 3’ scRNA-seq, we detected only 0.1% of reads assigned to gRNA from the library, whereas the 5’ scRNA-seq gave rise to 52.9% of reads assigned to gRNA. This is consistent with reliable assignment of the gRNA to the cell barcodes and high detection of gRNA levels from 5’ scRNA-seq (**Fig. S4B-G**) (see Methods). Moreover, in the 5’ scRNA-seq we could observe a threshold to separate cells with high gRNA expression from those with low gRNA expression, likely separating the true expression from the ambient, spurious, or background low-level detection (**Fig. S4G**). Using *in vivo* AAV Perturb-seq, we reliably detected gRNA identity in 63-83% of cells by 5’ scRNA-seq, which was about 7-10-fold higher than those by 3’ chemistry at 7%-14% (**Fig. 4D, Table S7**).

### Efficient cell labeling, high gRNA recovery, and validated CRISPR effects *in vivo*

Finally, we applied the AAV-based Perturb-seq system to a proof-of-principle *in vivo* screen targeting transcription factors with roles in brain development: *Foxg1*, *Nr2f1 (COUP-TFI)*, *Tbr1*, and *Tcf4*. Haploinsufficiency of these genes is associated with neurodevelopmental disorders and diseases, including Bosch-Boonstra-Schaaf optic atrophy syndrome (*Nr2f1*), Pitt-Hopkins Syndrome (*Tcf4*), FOXG1 syndrome (*Foxg1*) and TBR1 syndrome (*Tbr1*). These transcription factors play critical roles in cortical neuronal differentiation and progenitor maintenance, regional patterning, neuronal migration, and circuit assembly through the regulation of networks of other transcription factors (Chen et al., 2021; Greig *et al*., 2013; Hou et al., 2020). Most of their expression spans from embryonic to postnatal stages: *Tcf4* and *Nr2f1* are broadly expressed in most cell types, whereas *Foxg1* and *Tbr1* expression is restricted to subclasses of neurons including deep layer projection neurons and immature neurons (**Fig. S6A-B**) (Di Bella *et al*., 2021; La Manno *et al*., 2021).

**Figure 4.**
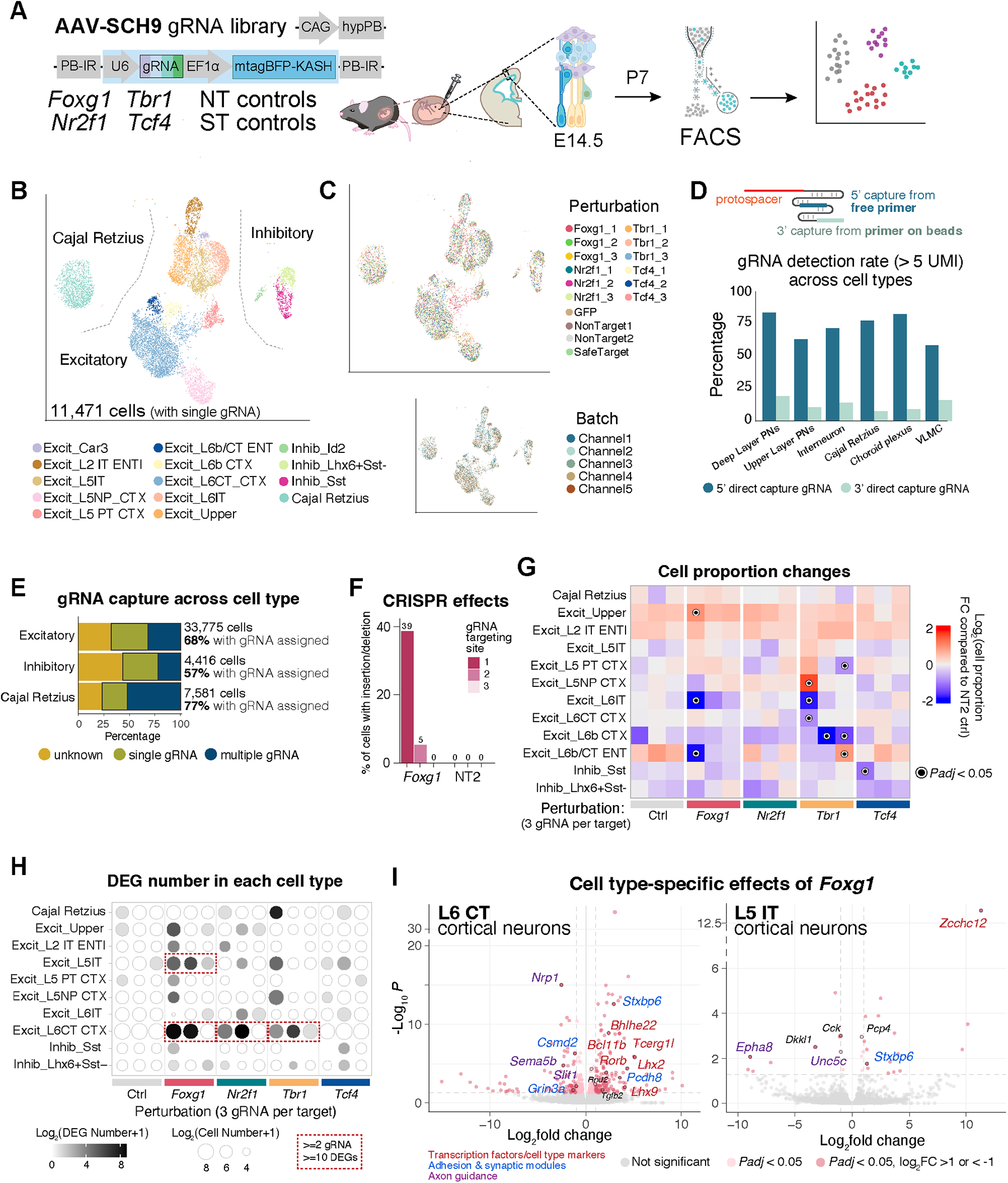
*In vivo* Perturb-seq identified cell type-specific changes across perturbations of transcription factors. (A) Schematics of the screen design. Shaded box indicates transposon flanked by the inverted repeats (IR); NT and ST indicate non-targeting and safe-targeting controls, respectively. (B) UMAP plot of filtered cells with a single perturbation, with each cell colored by annotated cell type. (C) UMAP plot of filtered cells, with each cell colored by gRNA identity estimated by DemuxEM with down-sampling (top) and batch/channel (bottom). (D) Top: schematics of 5’ and 3’ scRNA-seq capture mechanism. Bottom: comparing gRNA capture rate by measuring percentage of cells in each cell type with gRNA UMI number greater than 5 in 5’ and 3’ scRNA-seq, with cell types on the X axis, percentage of cells assigned gRNA identity on the Y axis. VLMC: vascular and leptomeningeal cells; PN: projection neurons. (E) Bar plot showing percentage of cells assigned to one or more gRNA across major cell types, color represents gRNA identity assignment. (F) Percentage of cells with insertion/deletion in *Foxg1* gRNAs targeting loci by *Foxg1* perturbation comparing to Non-Targeting control 2 (NT2) controls, extracted from scRNA-seq data. This plot only considers cells with at least one read overlapping the targeted region. (G) Heatmap showing cell type proportion changes by each perturbation. Color represents the log of cell proportion fold change compared to NT2 control; black rings highlight FDR adjusted *P*-value<0.05. (H) Dot plot of number of differentially expressed genes (DEG) (color) and cell number (size of dots) across cell type-perturbation combinations. Red boxes highlight the robust changes of <u>ý</u>10 DEGs supported by at least 2 gRNAs. (I) Volcano plots of cell-type specific effects on differential expressed genes of *Foxg1* gRNA1 perturbation in Layer 6 CT and Layer 5 IT excitatory neurons. Red and pink dots label significantly altered DEGs; black rings label the highlighted representative DEGs with their annotated functions: transcription factors and cell fate markers in red texts, adhesion and synaptic module genes in blue text and axon guidance-related genes in purple texts.

We designed four gRNAs for each gene by targeting coding exons closer to the 5’ end of the transcripts and tested them *in vitro* to select the three best-performing gRNAs to be included in the pool (**Fig. S4H, Table S2**). We pooled them equally along with four control gRNAs including non-targeting (NT) control and safe-targeting (ST) control gRNAs (Morgens et al., 2017) (**Fig. S4I**). AAV-SCH9 vectors expressing hypPB and pooled gRNAs were administered into embryonic lateral ventricles at E14.5 (**Fig. 4A**). At P7, the isocortex and hippocampal formation were micro-dissected and dissociated; the BFP-expressing cells were enriched and processed for droplet-based 5’ scRNA-seq with direct capture of gRNA (Replogle *et al*., 2020). In this experiment, we intentionally controlled the AAV injection titer to achieve <2% of cells transduced in the neocortex to limit multiple perturbation events (**Fig. S4J**), although this ratio can be further increased if testing combinations of perturbation is desired. Notably, with this intentional dilution, our system already achieves the collection of >10-fold more BFP^+^ perturbed cells than the conventional lentiviral labeling (**Fig. S4K**), which greatly streamlines the experiment.

Across five replicates (10x Chromium channels) with a total of 11 animals from two litters, we obtained a total of 50,075 cells profiled after the primary quality control (**Table S3**) (see Methods). Remarkably, this is much more efficient than our previous lentiviral-based work (to collect 46,770 cells over 17 litters and 163 animals) (Jin *et al*., 2020), demonstrating the scalability of this new platform. The cells were classified into six general cell types and 21 subclusters, annotated by direct comparison with public scRNA-seq data (see Methods; **Fig. S5A, Table S8**) (Yao et al., 2021).

To accurately assign the perturbation identities to cells, we evaluated multiple methods (see Methods). Based on this analysis, we decided on an approach involving first down-sampling the gRNA counts in each cell to minimize biases due to count differences (**Fig. S5G**), followed by DemuxEM, a tool designed for demultiplexing hashing barcodes (Gaublomme et al., 2019) (see Methods). We identified 16,067 cells with a single gRNA perturbation assigned, 16,373 cells with multiple gRNAs assigned, and 17,635 cells with no gRNA assigned, most of which were glia (**Fig. S5A, D**). Glial populations were associated with low gRNA recovery and low BFP detection (expressed from the AAV transgene), consistent with our previous characterization of AAV-SCH9 tropism (**Fig. 2, S5B, S5E**). Overall, we successfully assigned 57-77% of cells to perturbations in our data from the non-glia cell types, with a median of 152 UMI gRNA detected per cell and 9,687 UMI endogenous transcripts per cell in excitatory neurons (**Fig. 4E, S5E-F**). We further filtered low-quality cells with low numbers of UMI or a high percentage of intronic reads, which likely indicates they were cytoplasmic, nuclear debris, or a similar population (La Manno *et al*., 2021) (**Fig. S5C**).

Next, we focused our downstream analysis on 11,471 high-quality cells, each with only a single gRNA perturbation. We retained 14 annotated cell types, following the classification taxonomy of isocortex and hippocampal formation previously described (Yao *et al*., 2021). This included ten clusters of excitatory glutamatergic neurons including the L2-4 upper layer, L5/6 IT (intratelencephalic), Car3, PT (pyramidal tract), NP (near-projecting), CT (corticothalamic), and L6b; three clusters of GABAergic inhibitory neurons (Id2, Sst, and Lhx6+Sst-), and Cajal-Retzius cells (**Fig. 4B**) from a total of 18 subclusters **(Fig. S6C).** We did not observe any obvious batch effects on cell type across the five channels in UMAP space (**Fig. 4C**). Each gRNA had 318-1,065 cells assigned to it, giving us 1,609-2,527 cells perturbed for each gene and 3,316 control cells (**Fig. S6D**).

To evaluate the CRISPR induced loss-of-function and target gene dosage *in vivo*, we extracted the endogenous transcript reads from cells receiving a gRNA perturbation in the 5’ scRNA-seq libraries. We detected cells harboring insertion or deletion events within each gRNA-targeting region (**Fig. 4F, S6E, Table S8**): for example, among cells with at least one read in the associated target region, gRNA1 of *Foxg1* induced several distinct 1-base pair insertions in 39% of the cells, frameshift mutations leading to premature transcriptional termination downstream (**Fig. S6F**). This is likely an underestimation of the perturbation effects, as nonsense mediated decay could degrade much of the mutated mRNA. We speculate that different gRNAs, targeting the same gene, may yield variable phenotypic outcomes due to their differential efficacies in gene editing, introducing potential phenotypic heterogeneity. We also observed that wild-type transcripts were detectable in our data, suggesting that *in vivo* Perturb-seq could allow the analysis of the effects of both full knockout and heterozygous loss-of-function.

### Distinct cell type proportion changes by transcription factor perturbation *in vivo*

To analyze the cell type proportion changes resulting from each perturbation, we performed statistical tests considering the proportion differences across batches and gRNA identities, compared to a non-targeting control group (NT-2) (Phipson et al., 2022) (**Fig. 4G, S6G**). We decided to perform this analysis at the gRNA level, rather than the target-gene level, due to the known variability of gRNA performance (**Fig. 4F, S6E**). We also included other control gRNA groups, compared to this chosen control group NT-2, to test the robustness of the analysis, as we expected little to no effect in these control-to-control comparisons.

Different gRNAs targeting the same gene often showed a similar trend of changes, even if they did not all reach the same level of statistical significance (**Fig. 4G**). Notably, several neuronal classes showed changes of proportion by perturbations of *Foxg1*, *Tbr1* and *Tcf4*. As the most profound effect, cells perturbed by *Foxg1* (gRNA1) had a significant 9.9-fold reduction of L6 IT neurons, accompanied by a 2-fold increase of upper layer projection neurons (**Fig. 4G, S6H**). But this perturbation did not significantly impact the proportion of L6 CT neurons, which reside in the same laminar layer as L6 IT neurons. This result highlights the cell type-dependent effects of *Foxg1* in governing Layer 6 neuronal cell fate, with different effects in two neuronal classes within the same layer and sharing the same developmental lineage.

Intriguingly, we observed that *Tbr1* (gRNA1) perturbation, which had the highest efficiency among the three *Tbr1* gRNAs *in vivo*, is associated with a significant reduction in the proportion of deep layer excitatory neurons including L6 CT (1.9-fold) and L6 IT (3.6-fold). This is consistent with the known role of *Tbr1* in maintaining L6 neuronal identity (Bedogni et al., 2010; Fazel Darbandi et al., 2018), though our analysis provides a refined annotation of its cell type-specific effects. Moreover, this *Tbr1* perturbation led to a 3.1-fold increase in L5 NP excitatory neurons, a distinct effect from its role in L6 (**Fig. S6G-H**).

### Cellular context-dependent perturbation impact on deep layer glutamatergic neurons

To further explore the impact of each perturbation at the molecular level, we performed differential expression (DE) analysis for each perturbation (gRNA) across cell types. For each cell type, cells containing the same gRNA were compared to those with a control gRNA (NT-2). We included only the perturbation-cell type pairs with >50 cells, filtered out lowly expressed genes, and performed DE analysis with edgeR (McCarthy et al., 2012), exercising caution as recommended by existing literature (Soneson and Robinson, 2018) (see Methods).

First, we examined the number of significant DE genes across cell type-perturbation combinations (**Fig. 4H**). Control groups, when compared to the NT-2 control, showed very few to none (0-1) significant DE genes, as expected. Four perturbation-cell type combinations showed markedly altered gene expression (at least two gRNAs showed <u>≥</u>10 significant DE genes): *Foxg1* in L5 IT and L6 CT neurons; *Nr2f1* in L6 CT neurons; and *Tbr1* in L6 CT neurons. Some of these changes aligned closely with established roles described in the literature: *Foxg1* in enforcing L6 CT neuronal identity (Liu et al., 2022b); *Tbr1* loss-of-function causing defects in L6 CT neurons that acquire both L5- and L6-like identity and electrophysiological properties (Bedogni *et al*., 2010; Fazel Darbandi *et al*., 2018).

In late neurogenesis, *Foxg1* is critical for cell fate specification across neuronal cell types through regulation of transcription factor networks to maintain L6 neuronal identity (Liu *et al*., 2022b). Indeed, we found *Foxg1* (gRNA1) loss-of-function affected several signaling pathways in L6 CT neurons including expression changes in genes encoding adhesion and synaptic proteins (*Stxbp6, Grin3a*), axon guidance proteins (*Nrp1*, *Sema5b*), and transcription factors (*Bhlhe22*, *Lhx2*) (**Fig. 4I**). We identified upregulation of neuronal markers that are normally expressed in nearby layers, including L4 spiny stellate neuron marker *Rorb4* (4-fold increase) and L5 subcerebral/corticospinal projection neuron markers *Bcl11b* and *Tcerg1l* (1.5-fold and 34-fold increase) in these L6 CT perturbed neurons (**Fig. 4I**). These data suggest that, in L6 CT postmitotic neurons, *Foxg1* maintains the identity likely by actively suppressing other transcription factors and alternative cell fates; and its loss of function leads to increased expression of this network.

Additionally, we detected a sub-cluster of L6 CT neurons (cluster 7 in **Fig. S6C**), which was predominantly comprised of cells carrying *Foxg1* perturbations (**Fig. 4B, S6J, Table S8**). This subcluster expressed some typical cell type- or layer-specific markers (*Tcerg1l, Lhx2*), as well as markers unique to this cluster (*Nkd1*), suggesting this is not simply a cluster of doublets or empty droplets and is indeed likely to be cells with hybrid fates (**Fig. S6J**). The proportion analysis showed that *Foxg1* perturbation significantly increased the production of cluster 7, a strong effect that stood out as statistically significant across all three independent gRNAs (2-24-fold increase, *P-adj*<0.05) (**Fig. S6G, J**). The misregulation of transcription factor expression as well as the emergence of a hybrid neuronal subcluster (**Fig. S6J**), altogether, provided a high-resolution characterization of the altered cell fates when *Foxg1* failed to maintain L6 CT identity.

Importantly, we uncovered distinct patterns of gene regulation of *Foxg1* across cell types: the same perturbation induced different DE genes in different cell types and had little overlap (**Fig. S6K**). Amongst all the *Foxg1* (gRNA1) perturbation DE genes in the four deep layer neuron cell types: *Stxbp6* (Syntaxin binding protein 6) expression was upregulated by the perturbation in L5 PT, L5 IT, L5 NP, and L6 CT neurons (**Fig. 4I, S6I, K**). However, we found that there were more distinct, than shared, regulatory networks in different deep layer excitatory neuron types. Most of the *Foxg1*-knockout induced changes of transcription factors were highly specific to the cell types. For example, the DE genes observed in L6 CT were largely absent in L5 IT, L5 PT or L5 NP cell types (**Fig. 4I, S6I, K**). Additionally, it is known that *Foxg1-Lhx2* interactions are crucial for cortical hem formation; loss of *Foxg1* resulted in a reduction of *Lhx2* expression in progenitors during early embryogenesis at E9.5 (Chou and Tole, 2019). However, in perinatal and early postnatal stage, the time window of our analysis, *Lhx2* expression was substantially increased in the postmitotic L6 CT and L5 NP neurons following *Foxg1* perturbation (20 and 74-fold, respectively), and not significantly changed in L5 IT or L5 PT neurons. Since our perturbation agent AAV was administered at E14.5 in this experiment, which is days after L6 neurons were born, the effect is equivalent to a conditional knockout in the postmitotic neurons, rather than knockout in the progenitors and their previously reported effects in E9.5. These discoveries reveal *Foxg1*’s pleotropic effects on gene regulation that is spatiotemporally dynamic – with distinct molecular consequences in different cell types and developmental time windows. This also highlights the flexibility of this Perturb-seq platform to probe gene function over the course of development.

Taken together, our analysis uncovered the cell type- and developmental stage-specific role of *Foxg1* in maintaining L6 CT neuronal properties, through actively repressing alternative cell fates in postmitotic neurons. *Foxg1* loss-of-function led to the upregulation of several transcription factors which ultimately leads to hybrid cell identities as well as altered capacity for synaptogenesis and circuit assembly, distinct from its other known roles in the progenitors. These analyses, with high temporal, spatial, and cell type specificity, altogether, reveal key molecular pathways through which *Foxg1* orchestrates the cell fate determination and maturation of diverse neuronal cell types. Our data, altogether, demonstrate the potential and massively parallelizable capacity of the *in vivo* Perturb-seq platform to expand the scale and depth of genetic analysis of the highly cell type-specific regulatory networks from intact tissues.

## Discussion

### Perturb-seq platform with a modular design

Methods to target and manipulate mammalian cell types *in vivo* have been revolutionized in the past decade. Building on these efforts, a key advantage of this massively parallel *in vivo* Perturb-seq platform is its modularity: diverse serotypes and capsids of AAV can allow cell type- and tissue-specific targeting of even rare cell populations *in vivo*, expanding Perturb-seq from rodents to additional animal models (Chuapoco et al., 2023; Jang et al., 2023; Tabebordbar et al., 2021). However, AAV transgene expression could be diluted or eventually lost, depending on the cell division and turnover rate *in vivo*. Here, we demonstrated a versatile platform through an extensive screen of 86 serotypes across AAV phylogeny (**Fig. 1**) and incorporated a transposon system to stably integrate the transgene into the genome; both significantly enhanced the expression. We successfully tested the transposon system with three serotypes, AAV-SCH9, AAV9-PHP.eB and AAV9-PHP.S, for enhanced expression in central and peripheral nervous systems (**Fig. 3**). Our finding that AAV-SCH9 enabled rapid, onset within 48 hours, and sustained expression *in vivo* presents new possibilities for efficient manipulations in the dynamic development context. AAV-induced toxicity, which largely depends on the concentration and the cis-regulatory sequences used (Xiong et al., 2019), should be carefully evaluated to minimize confounding effects (Johnston et al., 2021). Future improvements to this platform could involve the use of alternative transposases, such as sleeping beauty (Querques et al., 2019; Ye et al., 2023; Ye et al., 2019), and precisely integrating transgenes into safe harbor loci (Hayashi et al., 2020; Pablo Perez-Pinera, 2011). These new integration systems could be modularly adapted and iteratively improved to be incorporated into Perturb-seq.

In the scRNA-seq data, we identified mutations in transcripts of several gRNA targets. Transcript genotyping may help differentiate between heterozygous and homozygous mutations in each cell to better inform phenotypic analysis (Nam et al., 2019), as most neurodevelopmental disorders risk genes are impacted by *de novo* heterozygous alleles. In addition, we can now directly capture gRNA from scRNA-seq dial-out readouts, rather than rely on distally located barcodes with the risk of recombination-mediated swapping (Xie et al., 2018). The tissue dissociation and single-cell isolation process may remove the less fit cells, which could introduce biases; in the future, *in vivo* Perturb-seq could be combined with multi-modal phenotyping, including *in situ* sequencing and spatial genomics (to identify the cellular migration, anatomical position, and morphology without tissue disruption) and single nucleus-based epigenetic profiling (to build cell state regulome networks) (Chong et al., 2022; Dhainaut et al., 2022; Feldman et al., 2019; Fleck et al., 2022; Mimitou et al., 2019; Rubin et al., 2019).

Similarly, it can also be coupled with additional perturbation tools such as CRISPRi, CRISPRa, CRISPRoff, or cDNA overexpression (Gilbert et al., 2014; Joung et al., 2023; Liu et al., 2022a; Nunez et al., 2021). These new features can be readily adopted in the next generations of AAV Perturb-seq as modular factors rather than relying on generating new mouse lines which can be costly and time-consuming.

### Identification of gene networks in cell fate determination during corticogenesis

Cortical function relies on diverse groups of specialized neurons that are born during embryonic development, orchestrated by networks of transcription factors and chromatin modifiers. Haploinsufficiency of many transcription factors has been confidently associated with human diseases and disorders. *In vivo* genetic screens are emerging platforms to interrogate gene function in the native cellular environment. We demonstrated the efficiency of our platform in capturing phenotypes that are tightly regulated spatiotemporally across diverse cell types (Klingler et al., 2021).

Through single cell-level analysis, we identified the role of *Foxg1* in governing Layer 6 neuronal cell fate, with different effects in different neuronal populations located in the same layer and sharing the same developmental lineage (**Fig. 4**). *Foxg1* perturbation decreased the proportion of L6 IT neurons; but the proportion of L6 CT neurons remains largely unchanged. In addition, the perturbation strongly altered the L6 CT neuronal cell fate, with the emergence of a hybrid cell state with upregulation of distinct patterns of transcription factors and cell adhesion molecules. Since the perturbation was introduced in late embryogenesis within the postmitotic neurons, we characterized the role of *Foxg1* in newborn neurons to sustain and maintain neuronal identity through actively repressing alternative cell fates, distinct from its role in regulating neuronal identities in the progenitors. These discoveries revealed *Foxg1*’s spatiotemporally dynamic role in cortical development – with distinct molecular consequences in different cell types and developmental time windows.

Amongst the *Foxg1*-knockout induced DE genes, only one gene (*Stxbp6*) was shared across the four cell types of deep layer glutamatergic excitatory neurons. The significantly altered transcription factor networks in L6 CT neurons are largely not changed in their neighboring neuronal cell types, showing highly restricted, cell type-specific effects of perturbations (West and Greenberg, 2011; Wong et al., 2018) (**Fig. 4, S6**). These biological discoveries required high-resolution and single cell-based phenotypic measurements to sample each of the many cell types. It is particularly worth noting that the functional activity of many risk genes’ actions depends on the cellular state, e.g. often being activity-dependent and sensitive to the environment (Boulting et al., 2021; Chen et al., 2019; Cheroni et al., 2020; Sanchez-Priego et al., 2022; Yim et al., 2020). Moreover, performing the experiments *in vivo* has the benefit that the system retains the endogenous cell-cell contacts, dynamic morphogen gradients, and the physiological environment – a key to decoding the relevant phenotypic changes that best resemble the disorder pathology.

### Towards in vivo, high-resolution phenotypic screens at scale

CRISPR screens, at variable scales, have been widely adopted across fields to study gene functions and to form new hypotheses (Przybyla and Gilbert, 2022). Most phenotypic readouts measure gRNA abundance as a proxy for proliferation or depletion; but biological complexity often transcends the ‘live or die’ phenotypes. Perturb-seq elevates the phenotypic resolution to the transcriptomic level with cell type specificity. Previously, *in vivo* Perturb-seq was limited by the number of cells that can be labeled and harvested from tissues, and the ability to capture gRNA reliably. Here, our AAV-based Perturb-seq achieved 10-fold higher labeling in embryonic brains (two batches for 50,075 cells using AAV, versus 17 batches 46,770 cells using lentiviral vectors) (Jin *et al*., 2020), with the ability to perturb >6% of cells in the adult brain with appropriate AAV vectors, in contrast to lentiviral vectors (<0.1%).

One of the key aims in functional genomics is to define the sites of actions for human disease and disorder risk genes: amongst all the heterogenous cell populations, including the central and peripheral nervous systems, what are the most vulnerable cell types and molecular networks that are affected by each genetic variant? Moreover, what are the cell type-and developmental time window-dependent functions of each gene? Towards this overarching goal, we designed our *in vivo* platform to be highly parallelizable and scalable, which is a required feature to systematically assay *hundreds* of risk genes across *hundreds* of cell types in diverse brain regions, with reasonable amounts of resources. Furthermore, this platform broadens perturbation and high-resolution screening possibilities to encompass a wider range of cell types and organs *in vivo*, such as Purkinje cells in the cerebellum, the peripheral dorsal root ganglia neurons, and potentially the enteric nervous system. We have demonstrated the efficiency and applicability of Perturb-seq in perinatal and adult tissues through several viral strategies, which makes our technology amenable to study other cellular contexts and disease mechanisms, including neurodegenerative diseases and aging. Emerging genomic tools will continue to enable us to move from associative observations toward understanding their relevance and function through perturbing panels of genetic variants to probe their cellular mechanisms and organ-scale impact on tissue development, homeostasis, and aging.

## Acknowledgement

We thank J. Dougherty and R. Mitra for sharing their experience working with hypPB; B. Li for advice on the use of DemuxEM; L. Tan for advice on the transposon insertion analysis; Y. Zhang for helping with the dorsal root ganglia dissection; Department of Animal Resources, Genomics Core and Flow Cytometry Core facilities at Scripps Research, Viral core at the University of California Irvine Center for Neural Circuit Mapping and Sanford Burnham Prebys, and Genomic core at Sanford Burnham Prebys for technical assistance; N.C., E.M., N.Y., S.S., A.P., L.Y., M.H., C.B. for advice on the manuscript; and all members of the Jin lab for their help and support. X.Z. and Y.L. were supported by Dorris Scholar Award. BW was supported by Mark Pearson Endowed Fellowship. J.Z.L. was supported by the Stanley Center for Psychiatric Research at the Broad Institute of MIT and Harvard. X.J. and this work were supported by the Simons Foundation for Autism Research Initiative Collaboration on Sex Differences, National Institute of Health (R01HG012819), Impetus grant, One Mind Rising Star Award, Klingenstein-Simons Fellowship Award, G. Harold and Leila Y. Mathers Foundation, Donald E. and Delia B. Baxter Young Investigator Award, Larry L. Hillblom Foundation, Astera Institute, and James Fickle.

## Author contributions

X.Z. and X.J. conceived the project and designed the experiments with inputs from all authors. X.X., L.T., and Q.W. designed and generated the barcoded AAV library reagents; X.Z. performed the *in vivo* experiments and tissue analysis with the help of B.W., Y.L., G.S.C., and X.J.; X.Z., B.W. and G.S.C. performed the single-cell RNA-seq experiment; X.Z., S.K.S., and K.K performed the Perturb-seq analysis under the supervision of J.Z.L. and X.J.; J.P. and X.Z. performed the AAV barcode analysis and whole genome sequencing analysis; Z.W. performed the qPCR experiments; A.A. constructed the plasmids that were used in this work with the help of X.Z.; X.Z. and X.J. drafted the manuscript with input from all authors.

X.J. and X.Z. are co-inventors on *in vivo* AAV-based Perturb-seq and CRISPR inventions filed by Scripps Research relating to the work in this manuscript.

## Tables with titles and legends

Table 1. AAV serotypes lists and barcodes in the 86-AAV and 14-AAV libraries.

Table 2. Summaries of oligos, primers, and AAV vectors used in this study.

Table 3. Summaries of scRNA-seq experiments in this study.

Table 4. Cell type classification and differential expressed genes in 14-AAV library E16.5 scRNA-seq data, related to Fig 2.

Table 5. Quantification of immunofluorescence data in this study.

Table 6. Whole genome sequencing analysis of transposon insertion events.

Table 7. Cell type classification of 5’ and 3’ scRNA-seq data, related to Fig 4.

Table 8. Cell type classification and differentially expressed genes in Perturb-seq data, related to Fig 4.

## Supplementary Materials

All the plasmid and viral vectors generated in this study will be deposited with Addgene. Other unique/stable reagents generated in this study are available from the lead contact with an executed Materials Transfer Agreement.

The data generated in this study has been submitted to the Mendeley Data (DOI:10.17632/hvb39r62xw.1) and NCBI Gene Expression Omnibus (accession no. TBD, will be provided by Oct 13, 2023) in both raw and processed forms. The accession number will be shared as soon as it becomes available. The analysis pipeline is deposited in the GitHub repository (https://github.com/jinlabneurogenomics).

### C57BL/6J, Cas9, and CD-1 mice

All animal experiments were performed according to protocols approved by the Institutional Animal Care and Use Committees (IACUC) of The Scripps Research Institute. E15 to P9 mice of varying sex and weight were used in the scRNA-seq experiments and mice ranging from E15 to adult were used in the immunohistochemistry experiments. All mice were kept in standard conditions (a 12-h light/dark cycle with ad libitum access to food and water).

### HT22 and HEK293FT cell lines

Mammalian cell culture experiments were performed in the HT-22 mouse hippocampal neuronal cell line (Millipore Sigma, #SCC129) or HEK293FT cell line (Thermo Fisher Scientific, #R70007) grown in DMEM (Thermo Fisher Scientific, #11965092) with 25mM high glucose, 1mM sodium pyruvate and 4mM L-Glutamine (Thermo Fisher Scientific, #11995073), additionally supplemented with 1× penicillin–streptomycin (Thermo Fisher Scientific, #15140122), and 5-10% fetal bovine serum (Thermo Fisher Scientific, #16000069). HT-22 cells were maintained at confluency below 80% and HEK293FT cells were maintained at confluency below 90%.

### Method details

#### Mammalian cell culture and time lapse imaging

All transfections were performed with PEI (Polysciences, #24765-1) in 24-well plates unless otherwise noted. Cells were plated at approximately 50,000 cells per well 16–20 hours before transfection to ensure 50-60% confluency at the time of transfection. For each well on the plate, 500-800 ng of transfection plasmids were combined with OptiMEM I Reduced Serum Medium (Thermo Fisher, #31985070) with PEI to a total of 50 µl. This solution was added directly to the media dropwise.

Cells were transfected and incubated in an In Cell6000 Analyzer (GE Healthcare) at 37°C and 5% CO2 and imaged with a 10x air objective. Images were collected every 4 hours for 20 hours. Image file names were blinded, and cell numbers were counted; fluorescence was analyzed with a custom script that is available on GitHub: https://github.com/jinlabneurogenomics.

#### RT-qPCR

HEK293FT cells were washed once with PBS, followed by trypsinization with TrypLE Express Enzyme (Thermo Fisher Scientific #12604021) and resuspension in PBS supplemented with 0.04% BSA (NEB, #B9000S). Cells were FACS purified at 4°C and collected in Trizol (Thermo Fisher Scientific, #10296010) with 5,000-50,000 cells per sample. RNA was extracted using Zymo Direct-zol RNA MiniPrep isolation kit (Zymo Research, #R2052). RT-qPCR was performed with Maxima H Minus Reverse Transcriptase (Thermo Fisher Scientific, #EP0753) and Power SYBR Green PCR Master Mix (Applied Biosystems, #4367659) with an equal mixture of the two RT primers: oligo (dT)18 primer (Thermo Fisher Scientific, #SO131) and direct capture primer 5’-TTGCTAGGACCGGCCTTAAAGC-3’. The qPCR probes are listed in Supplemental **Table S2**. The 2-ΔΔCT method was used for the analysis of qPCR data and normalized to GAPDH expression.

#### AAV vector construction and production

Viral vectors and plasmids were constructed as previously reported (Jin *et al*., 2020). The backbone plasmid contains the human U6 promoter to express one gRNA, and the EF1α promoter to express a fluorescent protein conjugated to the nuclear membrane localized domain KASH. Cloning of the vectors was done individually and confirmed by Sanger sequencing. The gRNA designs were defined using the online tool at benchling.com and the full sequences of the gRNAs used in this work are listed in **Table S2**. AAV production and titration was performed by the viral vector core facility at Sanford Burnham Prebys and UCI Center for Neural Circuit Mapping viral core.

#### Viral vector administration

AAV or lentivirus (0.5-1.5 μL per embryo, with AAV titer at 1-9×10^12^ U/mL and lentiviral titer at 9×10^9^ U/mL) was administered *in utero* to the lateral ventricle at E13.5-17.5 in CD1, C57BL/6J or Cas9 transgenic mice (Jax#026179) (Platt et al., 2014) for immunohistochemistry analysis and scRNA-seq. Adult mice were injected retro-orbitally with AAV (50-100 μL and ∼1-4e^11^ viral genome per animal) and perfused for immunohistochemistry experiments and nuclei flow cytometry.

#### AAV library barcode extraction

At 24-or 48-hours post transduction, labeled HT-22 cells and mouse primary cortical cells were purified with FACS. Genomic DNA was extracted from approximately 3,000 purified cells by using QuickExtract DNA Extraction Solution (Lucigen, #NC0302740) following the manufacturer’s protocol. The AAV serotype library was lysed by DNase I digestion and Proteinase K digestion. PCRs with genomic DNA were performed with NEBNext High-Fidelity 2X PCR Master Mix (New England BioLabs, #M0541L) with the following primers:

5’-CTTTCCCTACACGACGCTCTTCCGATCTgacgagtcggatctcccttt-3’

5’-GACTGGAGTTCAGACGTGTGCTCTTCCGATCTgcgatgcaatttcctcattt-3’

Amplicons were amplified to include adaptors and sequenced on iSeq 1000 or MiSeq platforms (> 2 million reads per sample; Illumina). BCL files were converted to FASTQ files using bcl2fastq (Illumina).

#### Immunofluorescent staining of brain sections and whole-mount DRGs

Embryonic brains were directly harvested after decapitation and frozen immediately on dry ice in OCT. Continuous sets of 15-20μm tissue sections were prepared on a cryostat, followed by the fixation for 15 min with 4% paraformaldehyde in PBS on ice. Postnatal pups and adult mice were anesthetized and transcardially perfused with ice-cold PBS followed by ice-cold 4% paraformaldehyde in PBS. Dissected brains were postfixed overnight in 4% paraformaldehyde at 4 °C. Postnatal and adult brains were embedded in 2% agar and 60-100μm tissue sections were collected on a vibratome.

The slides with mounted tissue sections were washed 4 times with PBS with 0.3% TritonX-100 and incubated with blocking media (10% donkey serum (Sigma Aldrich, #S30-100ML) in 0.3% TritonX-100 with PBS) for 2 hours at room temperature, then incubated with primary antibodies in the blocking media overnight at 4°C. Slides were washed with PBS with 0.3% TritonX-100 4 times. Secondary antibodies were applied at a 1:1000 dilution in blocking media and incubated for 2 hours at room temperature. Slides were then washed 4 times with PBS with 0.3% TritonX-100 and incubated with DAPI for 10 mins before mounting with Antifade Mounting Medium (Vector Laboratories, #H-1700-10). All images were taken using a Nikon AX Confocal Microscope with a 10x air or 20x air objective. Cell numbers were quantified manually with counter in ImageJ.

The primary antibodies and dilutions were: Chicken anti-GFP antibody (ab16901, 1:500; Millipore), Rabbit anti-GFP antibody (A-11122, 1:500; Invitrogen), Rabbit anti-RFP (600-401-379, 1:500; Rockland), Rabbit anti-Tbr1 (ab31940, 1:500: Abcam), Rabbit anti-Tbr2 (ab183991, 1:500: Abcam), Rat anti-Ctip2 (ab18465, 1:1000, Abcam), Rabbit anti-Pax6 (Cat#901302, 1:500, BioLegend), Rabbit anti-HA tag (5017, 1:500; Cell Signaling), Rat anti-HA tag (11867423001, 1:500, Roche), Chicken anti-GFAP (ab4674, 1:500, Abcam) and Goat anti-Iba1 (ab5076, 1:500, Abcam).

Mouse dorsal root ganglia (DRGs) were extracted and post-fixed in 4% PFA for 1 hour before being washed with PBS. DRGs were mounted onto silicone isolators and mounted using EasyIndex.

#### Nuclei isolation and FACS-enrichment for genomic analysis

For the whole genome sequencing, >170,000 tdTomato^+^ nuclei were sorted using Sony cell sorter (SH800) into DNA/RNA Shield (Zymo Research, Cat# R1100-250) and processed for DNA purification using the Quick DNA Microprep Kit (Zymo Research, #D3020) to isolate >500 ng genomic DNA. Genomic DNA was sequenced to 60x coverage with paired-end (read 1 = 150 bases, read 2 = 150 bases) Illumina sequencing (Novogene).

#### Tissue Dissociation and FACS-enrichment for genomic analysis

Tissue dissociation was performed with the Papain Dissociation kit (Worthington, #LK003150) in a modification of a previously described protocol (Jin *et al*., 2020). Briefly, young mice were anesthetized then disinfected with 70% ethanol and decapitated. The brains were quickly extracted and gently dabbed with a PBS-soaked Kimwipe (Kimberly-Clark) to remove the meninge and fibroblasts. Cortices were micro-dissected in ice-cold dissection medium (Hibernate A medium (Thermo Fisher Scientific, #A1247501) with B27 supplement (Thermo Fisher Scientific, #17504044) and Trehalose (Sigma Aldrich, Cat# T9531) under a dissecting microscope. Microdissected cortices were transferred into papain solution with DNase in a cell culture dish and cut into small pieces with a razor blade. The dish was then placed onto a digital rocker in a cell culture incubator for 30 mins with rocking speed at 30 rpm at 37°C. The digested tissues were collected into a 15 mL tube and triturated with a 10 mL low bind plastic pipette 20 times and the cell suspension was carefully transferred to a new 15 mL tube. 2.7 mL of EBSS, 3 mL of reconstituted Worthington inhibitor solution, and DNase solution were added to the 15 mL tube and mixed gently. Cells were pelleted by centrifugation at 300 g for 5 mins at 4°C, followed by washing with 8ml cold dissection medium at 200g for 5 min at 4°C. Cells were resuspended in 0.5 mL ice-cold dissection medium with 10% fetal bovine serum (FBS) (Thermo Fisher Scientific, #16000069) and SYTOX dead cell stain (Invitrogen, #S34859), and subjected to FACS purification using Sony cell sorter (SH800).

After collection, the cells were immediately centrifuged and resuspended in ice-cold PBS with 0.04% BSA (NEB, #B9000S). Each 10x scRNA-seq library was prepared by combining the FACS sorted cells from 1-2 litters (5-8 animals) of E15 or P7-9 animals harvested on the same day (**Table S3**). We performed the dissociation, FACS purification and resuspension within 3 hours while keeping the cells on ice to prevent necrosis.

The scRNA-seq libraries were constructed using the Chromium Next GEM Single Cell 3’ Solution v3.1 kit with Feature Barcode Technology or the Chromium Next GEM Single Cell 5’ Solution v2 kit with Feature Barcode Technology (10x Genomics) following the manufacturer’s protocol. The gene expression library was sequenced with NextSeq500 high-output 75-cycle kits (Illumina) with sequencing saturation to ensure greater than 20,000 reads coverage per cell (R1: 26 bases, R2: 46 bases). The CRISPR gRNA screening library was sequenced with Illumina iSeq100 300-cycle (R1: 151 bases, R2: 151 bases) and Nextseq500 mid-output 150-cycle kits (Illumina) (R1: 73 bases, R2: 74 bases).

#### AAV barcode enrichment from scRNA-seq library

Following whole transcriptome amplification (WTA) in the 10x Chromium library construction, a fraction of the WTA product was used to amplify AAV serotype barcodes as well as cell barcodes using a dial-out PCR strategy (**Table S2**). Briefly, 10 ng of WTA libraries were amplified for 11 cycles of PCR with:

AAVlib-dialout-NGS1:

GACTGGAGTTCAGACGTGTGCTCTTCCGATCTGACGAGTCGGATCTCCCTTT

Read1-F:

CTACACGACGCTCTTCCGATCT

followed by a 1X SPRIselect beads cleanup (Beckman, #B23318). The sample was amplified another 24 cycles with TruSeq indexed primers and gel purified. The final dial-out library was sequenced along with transcriptome library with NextSeq500 high-output 75-cycle kits (Illumina) flow cell (R1: 26 bases, R2: 46 bases).

### Quantification and Statistical Analysis

All images were analyzed with ImageJ (NIH), Photoshop (Adobe) and Illustrator (Adobe). Cells were counted manually from blinded files using the ImageJ CellCount function.

#### AAV barcode analysis in the primary serotype screen

FASTQ files of Illumina libraries were mapped to AAV barcodes using a custom script (https://github.com/jinlabneurogenomics/aavperturbseq/fastq_barcodemapping.py). Briefly, sequences that began with the correct initial primer sequence were kept. Of these sequences, the barcode sequence following the initial primer sequence was compared to our list of AAV barcodes and assigned to matching AAV barcodes with Levenshtein distance less than 2.

The barcode counts matrix was analyzed using DESeq2 v1.40.2 (Love et al., 2014). The DESeqDataSetFromMatrix command was used to create the DESeq object with the library counts as the reference level. Results were tabulated for each condition using alpha=0.05 as the threshold for significance. Volcano plots were produced using the R package EnhancedVolcano v1.18.0 and heatmaps with pheatmap v1.0.12.

#### Transposon integration site analysis

A custom reference genome was created by appending the transposon reporter plasmid to the mm39 mouse genome as an additional chromosome. FASTQ files were aligned to the custom genome using bwa mem (v0.7.17) (Li and Durbin, 2010) using the SP5M flags. The resulting files were filtered for reads that aligned to the piggyBac plasmid and their pairs using samtools v1.15.1 (Li et al., 2009). After checking the distribution along the plasmid of these reads, they were further filtered down to reads aligning to the ends of the insert (position 1653-2053 and 6096-6496) within the plasmid. The filtered files were then parsed using pairtools v0.3.0 (Song et al., 2022) and default settings. They were then filtered for junctions (pair_type of UU, UR, or RU with one end aligning to the mouse genome) that were then validated manually.

To annotate the integration sites, the annotatePeak function from the R package ChIPseeker (v1.36.0) (Yu et al., 2015) was used with the TSS set as +/-3,000 bp. The R package circlize v0.4.15 (Gu et al., 2014) was used to create a circos plot to visualize integration sites.

#### scRNA-seq data processing

BCL files of transcriptome libraries were used to generate FASTQ files using the default parameters by “cellranger mkfastq” command (Cell Ranger v7.1.0) (Zheng et al., 2017). The gRNA library or AAV barcode dial-out library was demultiplexed using bcl2fastq. The “cellranger count” command was used to align the transcriptome reads to the mouse genome reference mm10 (GENCODE vM23/Ensembl 98) and generate a gene expression count matrix, using expect-cells = 9,000. The AAV barcodes or gRNA reads were quantified at the single-cell level with the feature-ref flag in Cell Ranger.

#### Cell type classification and cell identity annotation

For the AAV serotype secondary screen and comparing 5’ vs 3’ scRNA-seq, filtered count matrices from Cell Ranger were loaded into R v4.3.0 with the Read10X command from Seurat v4.3.0.9003 (Hao et al., 2021) and loaded into a Seurat object with CreateSeuratObject, filtering out cells with < 500 genes or mitochondrial count percent > 25%. The data were log normalized and the 2,000 most variable features were selected by FindVariableFeatures. In each analysis, two conditions (sort vs unsort, 5’ vs 3’) were integrated by the variable features agreed across datasets by IntegrateData. The integrated data was scaled by ScaleData and PCA was performed by RunPCA. A UMAP was generated with RunUMAP and clustering was performed by FindNeighbors and FindClusters (with default parameters, except for dims = 1:25, resolution = 0.3 for the sort vs unsort, or 0.2 for 5’ vs 3’). Clusters were assigned to cell types based on the known markers (Di Bella *et al*., 2021; La Manno *et al*., 2021; Tasic *et al*., 2018). In the AAV serotype screen, non-cortical cell clusters were removed, while cell clusters with too few cells (<150 cells) were removed in the 5p vs 3p analysis. The data were re-clustered (dims = 1:28, resolution = 0.3 for the AAV serotype screen library or 0.2 for 5’ vs 3’). The AAV barcode or gRNA identity was assigned for cells with >5 UMI detected.

#### Perturb-seq data processing: cell type classification, cell identity annotation, and perturbation identity annotation

The filtered count matrices from Cell Ranger were loaded into R v4.0.3 with the Read10X command from Seurat v4.0.0 (Hao et al., 2021) and loaded into a Seurat object with CreateSeuratObject, filtering out cells with <500 genes. The data was normalized with the NormalizeData command, with *normalization.method=“LogNormalize”* and *scale.factor=1,000,000* and variable genes were selected with FindVariableFeatures. Doublet scores were calculated with scds v1.6.0 (Bais and Kostka, 2020) separately for each 10x channel for use in cluster annotation (see below). The UMI count data for the variable genes were extracted and used as input to scGBM v0.1.0 (Phillip and Jeffrey, 2023) with M=20 and subset=30000, resulting in a 20-dimensional embedding of the count data that was added to the Seurat object. A UMAP was calculated on this reduction using RunUMAP, and clustering was performed on this reduction with FindNeighbors and FindClusters with otherwise default settings. The UMI count matrix for the gRNAs produced by Cell Ranger was also added to this Seurat object as an additional assay.

Quality control (QC) metrics for each channel were calculated with CellLevel_QC tool (https://github.com/seanken/CellLevel_QC) and loaded into the metadata for the Seurat object. The % mitochondrial reads was also calculated for each cell. Azimuth v0.3.2 was used to produce an initial annotation of the data using a single cell reference from the Allen brain atlas (Yao *et al*., 2021). Clusters with high percent intronic reads or high doublet scores were removed, as were cells with >20% intronic reads or >10% mitochondrial reads. Clusters were then labeled with cell type using the cell type labels from Azimuth and by comparing DE genes from our dataset to DE genes from the Allen brain atlas dataset (DE genes between clusters were calculated with presto v1.0.0) (Korsunsky et al., 2019).

We next built a Nextflow pipeline (DSL v2) (Di Tommaso et al., 2017) that took the Cell Ranger output, extracted the guide RNA data into a csv file, assigned cells to guides with DemuxEM (v0.1.5) (Gaublomme *et al*., 2019), and extracted the resulting perturbation labels in the form of a tsv file (where the creation of the tsv and csv was performed using pegasusio v0.2.11 (Li et al., 2020) for loading the data and pandas v1.2.4 for saving the data (pandas-development-team, 2023)). These tables were then loaded into the Seurat objects metadata. We also built a version of the pipeline that allowed for down sampling before running DemuxEM (the down sampling approached mentioned in the main text). In particular, for each 10x channel and each gRNA, if there were more than 50,000 UMIs assigned to that gRNA, the pipeline down-sampled the gRNA UMI counts for that gRNA using the binomial function from numpy.random (Harris et al., 2020), where the proportion used for down sampling was equal to 50,000 divided by the total number of UMI coming from the gRNA (resulting in an expected value of 50,000 UMIs per gRNA per channel after the down sampling). Similarly, we built a Nextflow pipeline to extract reads mapping to the BFP or GFP sequence. The pipeline took in the BAM file produced by Cell Ranger, extracted unmapped reads with samtools v1.8 (using the command “samtools view -f 4 -b”) (Li *et al*., 2009), extracts UMIs and CBCs with umi_tool v1.0.1 (Smith et al., 2017), mapped these reads to the sequences for GFP and BFP (including 5’ and 3’ UTR regions) with minimap2 v2.11 (using the arguments -ax sr) (Li, 2018), and transformed them into a BAM file with samtools view -b. Unmapped reads were discarded. For the remaining reads, the transcript that read was mapped to (GFP or BFP) was added to that read in the BAM file as an additional tag (XT tag) using the awk command. The resulting BAM file was sorted, indexed, and the number of UMIs mapping to GFP and BFP for each cell was extracted with command umi_tools count –per-gene –gene-tag=XT –per-cell. These counts were also loaded into the Seurat object.

In order to accurately assign the perturbation identities to cells, we compared three methods: the built-in gRNA assignments from Cell Ranger, a tool for demultiplexing the hashing barcodes known as DemuxEM (Gaublomme *et al*., 2019), and down sampling gRNA counts followed by DemuxEM. We found that all three methods performed well, though we saw evidence showing that the standard DemuxEM approach led to biases in cell type proportion due to large differences in the number of gRNA UMIs recovered per guide (**Fig. S5G**). For example, we observed a strong correlation between the average number of gRNA UMIs recovered from a given gRNA and the proportion of Cajal Retzius cells among cells assigned to that gRNA (Spearman correlation=0.84, *P-value*=2.934e-05); this effect disappeared after the down sampling (Spearman correlation=-0.03, *P-value*=0.89). Cell Ranger showed evidence of this bias but reported a larger than expected number of doublets, particularly in the Cajal Retzius cluster (with 71% of high-quality Cajal Retzius cells being assigned as doublets in Cell Ranger, versus 60% in the down sampling approach and 52% in the standard DemuxEM approach). We used the DemuxEM with down sampling for the full analysis.

Non-neuronal cells (cells not labeled as Excitatory, Inhibitory, or Cajal-Retzius neurons) were removed from the Seurat object, as were cells with < 3% intronic reads. Excitatory and Inhibitory cells with < 3,000 genes per cell were filtered out, as were Cajal-Retzius cells with < 2,000 genes per cell. Finally, all cells that were not assigned to exactly one guide by DemuxEM were removed as well. This Seurat object was used for downstream analysis.

The differential expression between clusters/cell types reported in the supplement (**Table S8**) was calculated with Seurat’s FindMarkers command.

#### Detecting insertions and deletions in targeted regions

For each targeted gene and each cell barcode, the reads overlapping that gene in each 10x channel were extracted from the Cell Ranger BAM files with samtools view and combined into one BAM file with samtools concat, which was sorted and indexed with samtools. The sinto filterbarcode command (Stuart et al., 2021) was used to split the BAM file into one BAM file for each perturbation, consisting of cells in the final analysis assigned to each gRNA (excluding cell barcodes that occur in multiple 10x channels). The resulting BAM files were indexed. This resulted in one BAM file for each targeted gene and each gRNA, consisting of reads in cells assigned to that gRNA/perturbation identity that overlap the targeted gene.

For each targeted region, we counted the number of reads overlapping that region for each perturbation, as well as the number of reads with an indel (insertion or deletion) in that region. This was performed with a pysam (https://github.com/pysam-developers/pysam) based python script. The script loaded each read one by one from the BAM file. For each position in each read it used get_reference_positions to: 1) check if that position was in the range required 2) if it was in the range, checked if that location was an insertion (excluding insertions at the beginning and end of reads to avoid soft clipped regions) 3) if it was in the range and did not contain an insertion, checked if that location was a deletion (if there was a difference of > 1 bp between the mapping position of the current base and the previous base). The results were then tabulated into a pandas data frame and returned before being saved then loaded into R for downstream analysis.

#### Perturbation-associated analysis

Having assigned a gRNA and a cell type to all the cells, we looked at cell type proportion changes across gRNAs. To do so, we selected a control group (Non-Targeting control 2) and compared each of the perturbation and control groups containing other gRNAs to this group. Statistics for these pairwise composition comparisons were computed using the propeller.ttest function from speckle (R package v0.99.7) (Phipson *et al*., 2022). The proportions were first transformed using arcsin square root transformation, and the batch (10x channel) was additionally considered as another fixed effect to the linear models. Cell types and clusters with less than 200 cells overall (Excit_Car3 and Inhib_Id2 for the cell type-level comparisons and clusters 5, 25 and 26 for the cluster-level comparisons) were excluded from this analysis. Results were collected and visualized together using ComplexHeatmap (R package v2.14.0) (Gu et al., 2016). For this visualization, effect sizes were capped at the absolute value of 2.

We next looked for genes that were differentially expressed in each gRNA and within each cell type. Using a control gRNA (Non-Targeting control 2), we compared each gRNA group population to the control. Statistics were computed with a pseudo-bulk approach using edgeR (R package v3.40.2) (McCarthy *et al*., 2012). Lowly expressed genes (<10 supporting reads and < 0.1 normalized expression) were identified within each comparison and excluded from the DE test. Additionally, we applied surrogate variable analysis (sva; R package v3.46.0) (Leek and Storey, 2007) to capture potential batch-or technical artifact-related variable from the data and add it to the model testing for DE (n.sv=1). Computed *P*-values were adjusted using the Benjamini-Hochberg method.

## Supplemental information titles and legends

**Figure S1.**
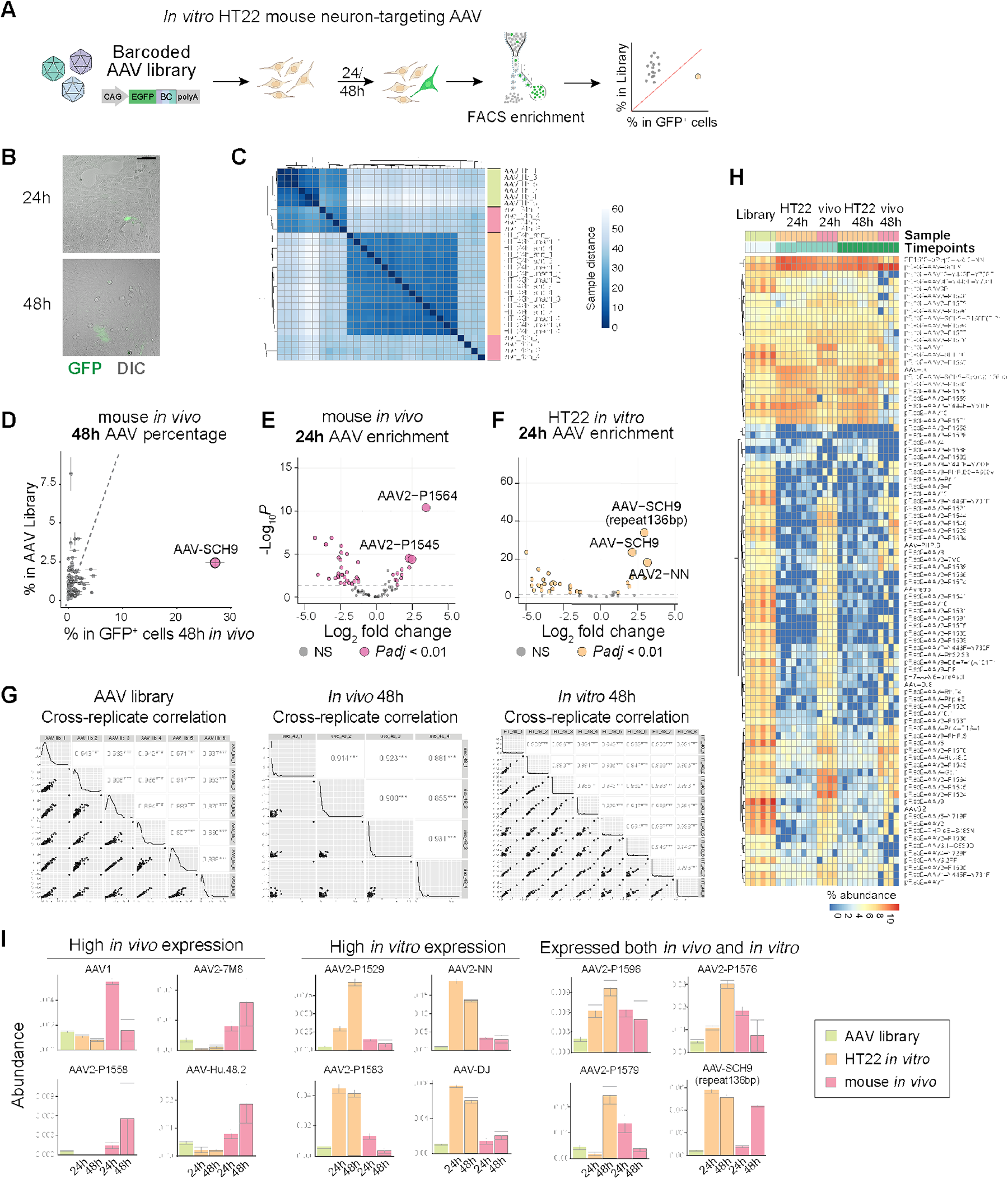
AAV serotype *in vitro* and *in vivo* screen reveals a candidate vector AAV-SCH9 to target newborn neurons in developing mouse brain. (A) Schematics of the *in vitro* AAV serotype screen: to introduce the pooled AAV library in HT22 cells followed by next generation sequencing of GFP^+^ cells to analyze the barcode. (B) Fluorescent and bright field images of HT22 24- or 48-hours post transduction. (C) Heatmap of sample correlation across the barcode distribution measurements from the initial AAV library, HT22 cells and mouse brain 24- or 48-hours post transduction. Each row represents a replicate library of the condition. Colors indicate pairwise Euclidean distance between each sample. (D) Scatter plot showing AAV serotype proportion in mouse brain 48 hours post transduction (X axis) compared to the initial AAV library (Y axis). (E-F) Volcano plot showing AAV serotype enrichment in mouse brain 24 hours post transduction (E) and in HT22 cells 24 hours post transduction (F). Highlighted serotypes were marked with bigger size dots with text labels. Dotted lines label log2FC = 1 or – 1. (G) Scatter plot comparing cross replicates of experiments and their correlations within the barcode abundance analysis of the initial AAV library, HT22 cells and mouse brain 48 hours post transduction. The values on the X and Y axes are the barcode abundance values. Numbers indicate Pearson correlation coefficient and asterisks indicate *P*-value <0.001. (H) Heatmap of the full 86 AAV serotype abundance in AAV library, HT22 cells and mouse brain 24- or 48-hours post transduction; each row represents the abundance of an AAV serotype. Color indicates the percentage abundance of each serotype in the sample. (I) Bar plots of representative AAV serotype barcode abundance in the initial AAV library, HT22 cells and mouse brain 24- or 48-hours post transduction. Error bars indicate standard error of the mean. Scale bar indicates 100 μm (in B).

**Figure S2.**
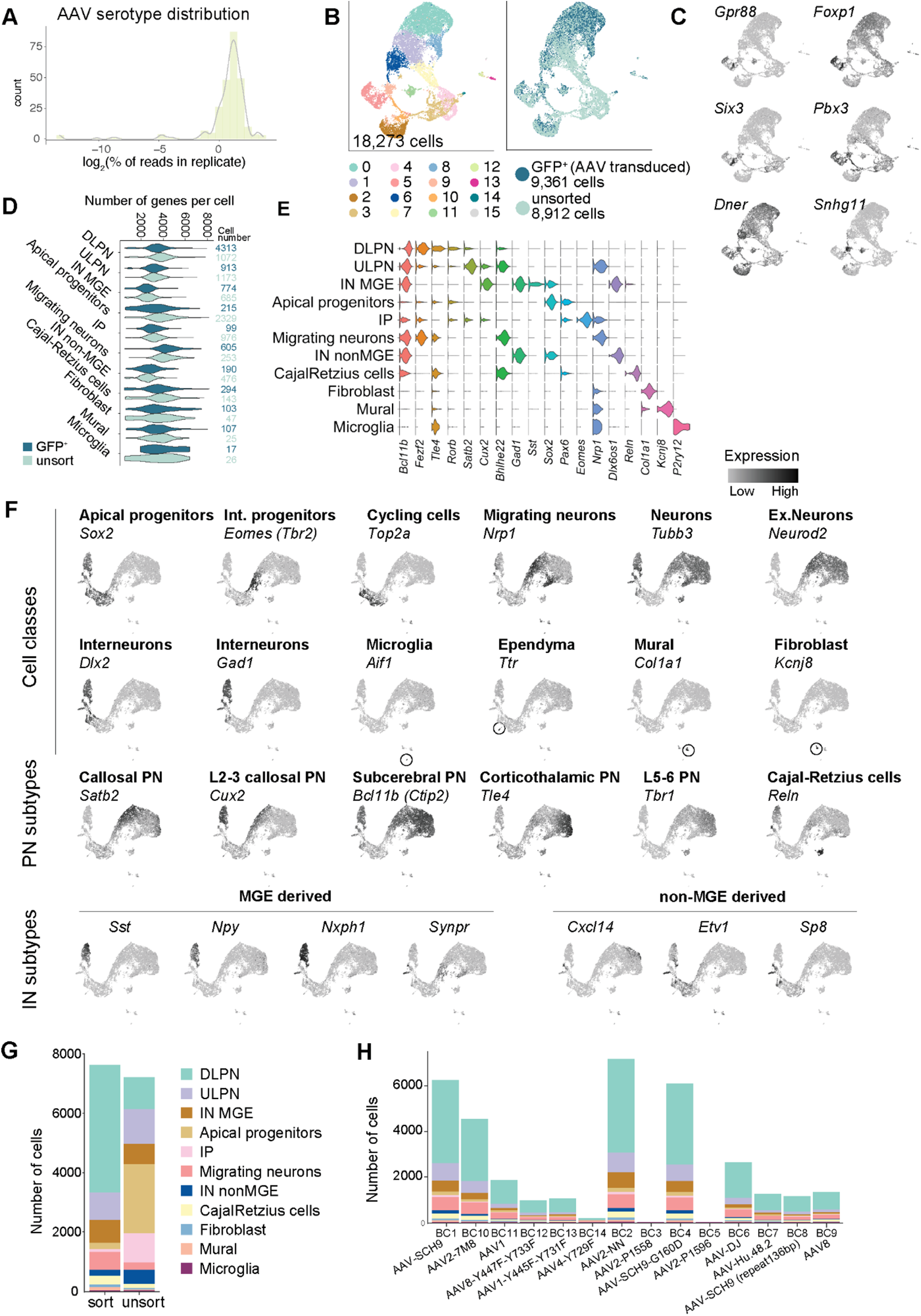
AAV-SCH9 tropism *in vivo* with single-cell resolution from intact tissues. (A) Histogram and density plot of serotype barcode distribution in the secondary 14-AAV library. (B) UMAP visualization of main clusters of transduced (GFP^+^) and total (unsorted) cell populations. (C) UMAP showing expression of non-cortical cell marker genes. (D) Violin plots of the number of genes detected per cell across diverse cell types. (E) Violin plots of canonical marker gene expressions across cell types annotated in the dataset. (F) UMAPs plots colored by marker genes for major cell types. (G) Bar plot of cell number distribution in each identified cell type across the transduced (sort) and total populations (unsort). (H) Bar plot of number of cells in each identified cell type, assigned to each of the 14 AAV barcodes. Y axis shows the number of cells for each cell type (sum of 3 separated barcodes for each serotype).

**Figure S3.**
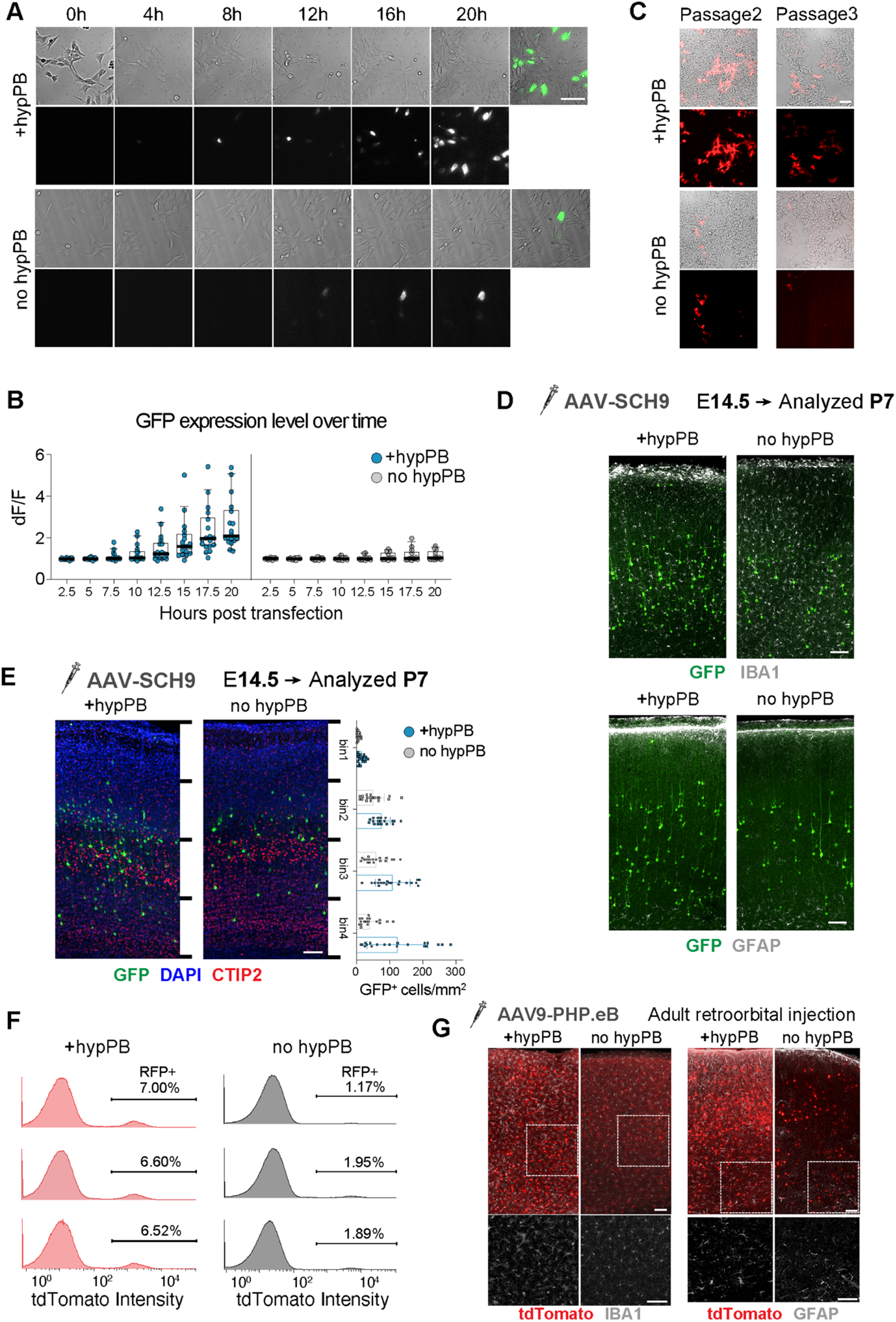
hypPB transposon effectively enhances and stabilizes expression in embryonic and adult brains. (A-B) Co-transfection of hypPB enhanced the expression of transgene *in vitro*. (A) Timelapse images of HT22 cells transfected with or without hypPB along with the reporter transposon expressing GFP. (B) Boxplots of quantification of the raw fluorescence level normalized to the background fluorescence (dF/F) showed elevated fluorescent signals in the presence of hypPB (left) relative to no hypPB (right). (C) Co-transfection of hypPB sustained transgene expression *in vitro* over 3 cell passages, without any selection. We performed the analysis, with and without hypPB, in HEK293 cells and followed by 1:10 passages every 3 days. (D) Embryonic co-transduction with AAV-SCH9-hypPB did not introduce substantial gliosis or cellular toxicity at P7. (E) Co-transduction with AAV-SCH9-hypPB increased GFP^+^ cells across cortical laminar layers, which were evenly divided into four bins. (F) Density plots of fluorescence signals from nuclei isolated from adult brains that were co-transduced with and without AAV9-PHP.eB-hypPB: hypPB increased tdTomato^+^ nuclei percentage. (G) Elevated gliosis, IBA1 and GFAP expression, with AAV-PHP.eB-hypPB transduction. Scale bars indicate 100 μm (in A, C, D, E and G).

**Figure S4.**
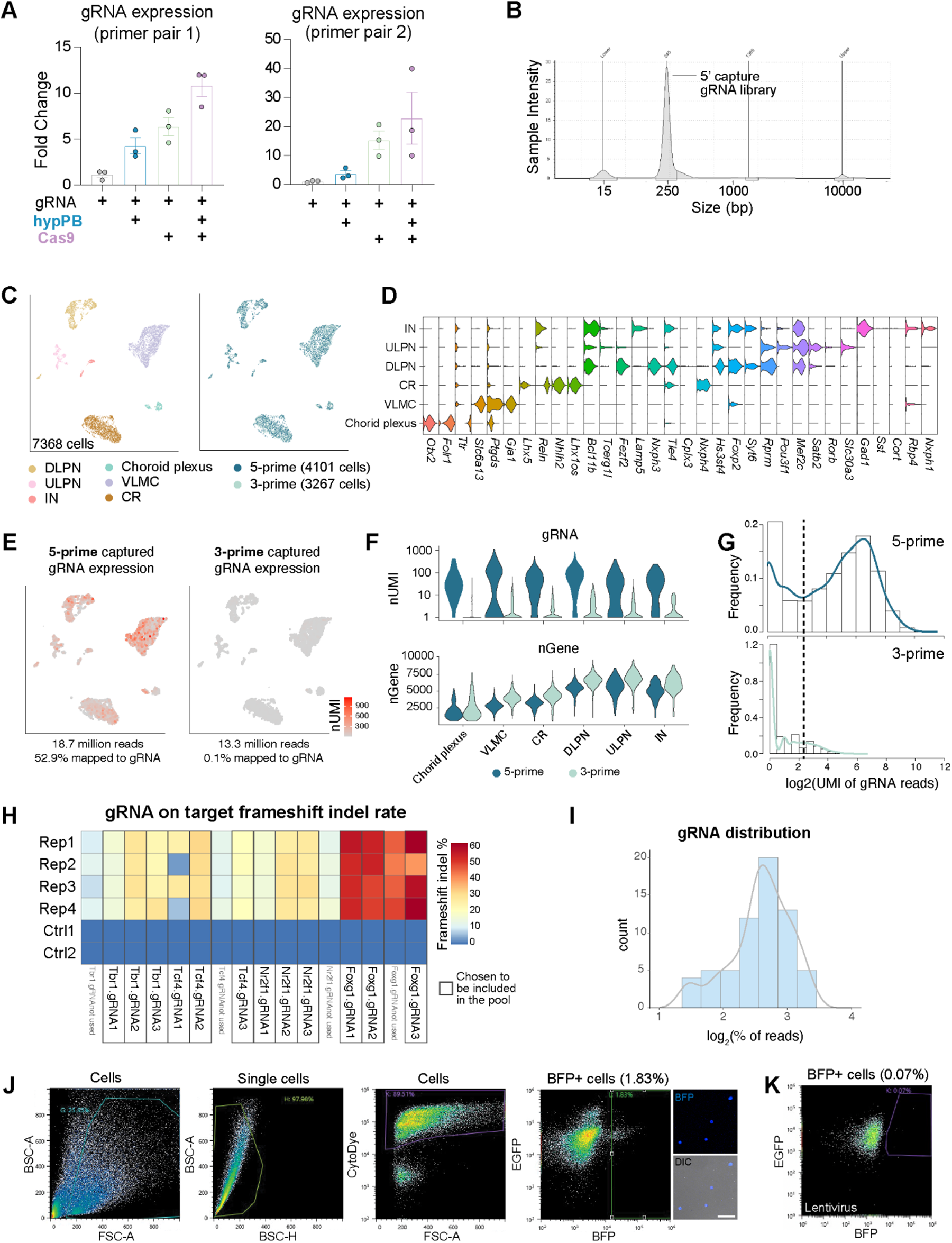
AAV-SCH9 and transposon system permits gRNA capture with sparse scRNA-seq readout. (A) qPCR analysis revealed both Cas9 and hypPB co-expression enhanced gRNA expression level, using two sets of independent primer sets (for the same target). The Y axis is fold change relative to *Gapdh*. Each point represents a replicate, error bars indicate standard error of the mean. (B) A representative Tapestation trace of the gRNA direct capture library from a 5’ 10x scRNA-seq library; a failed gRNA capture will result in no peak in the 250 bp region. (C) UMAP plots of neocortical cells from 3’ and 5’ scRNA-seq libraries; each cell is colored by annotated cell type (left) and 10x scRNA-seq technology (right). Cell types include upper and deep layer projection neurons (ULPN, DLPN), interneurons (IN), choroid plexus, Cajal-Retzius cells (CR) and vascular and leptomeningeal cells (VLMC). (D) Violin plots of canonical marker gene expression across cell types. (E) Feature plots of gRNA expression level detected in 5’ and 3’ kits: with comparable sequencing depth, only 0.1% were mapped to gRNA from the 3’ kit, whereas >50% were mapped in the 5’ kit. (F) Violin plots of transcriptomic and gRNA capture efficiency in 5’ and 3’ gRNA direct capture library preparation. Detected gRNA expression level per cell was overall higher in the 5’ kit, while the endogenous transcripts detected per cell were similar. (G) Histograms and density plots of gRNA expression level per cell in 5’ and 3’ gRNA direct capture kits. Dashed line represents a threshold to separate cells with high gRNA expression from those with low gRNA expression, likely separating true expression from the ambient, spurious expression. (H) Heatmap of gRNA on-target frameshift insertion or deletion (indel) rate. Boxes highlight the gRNAs chosen to be included in the *in vivo* screen. (I) Histogram of distribution of 16 gRNAs in AAV-SCH9 gRNA pool. (J) FACS enrichment gating for AAV-transduced cells (∼1.83% of total cells *in vivo*). Representative images of sorted, perturbed, BFP^+^ cells. (K) FACS enrichment gating for lentivirus transduced cells (∼0.07% of total cells *in vivo*). Scale bar indicates 50 μm (in J).

**Figure S5.**
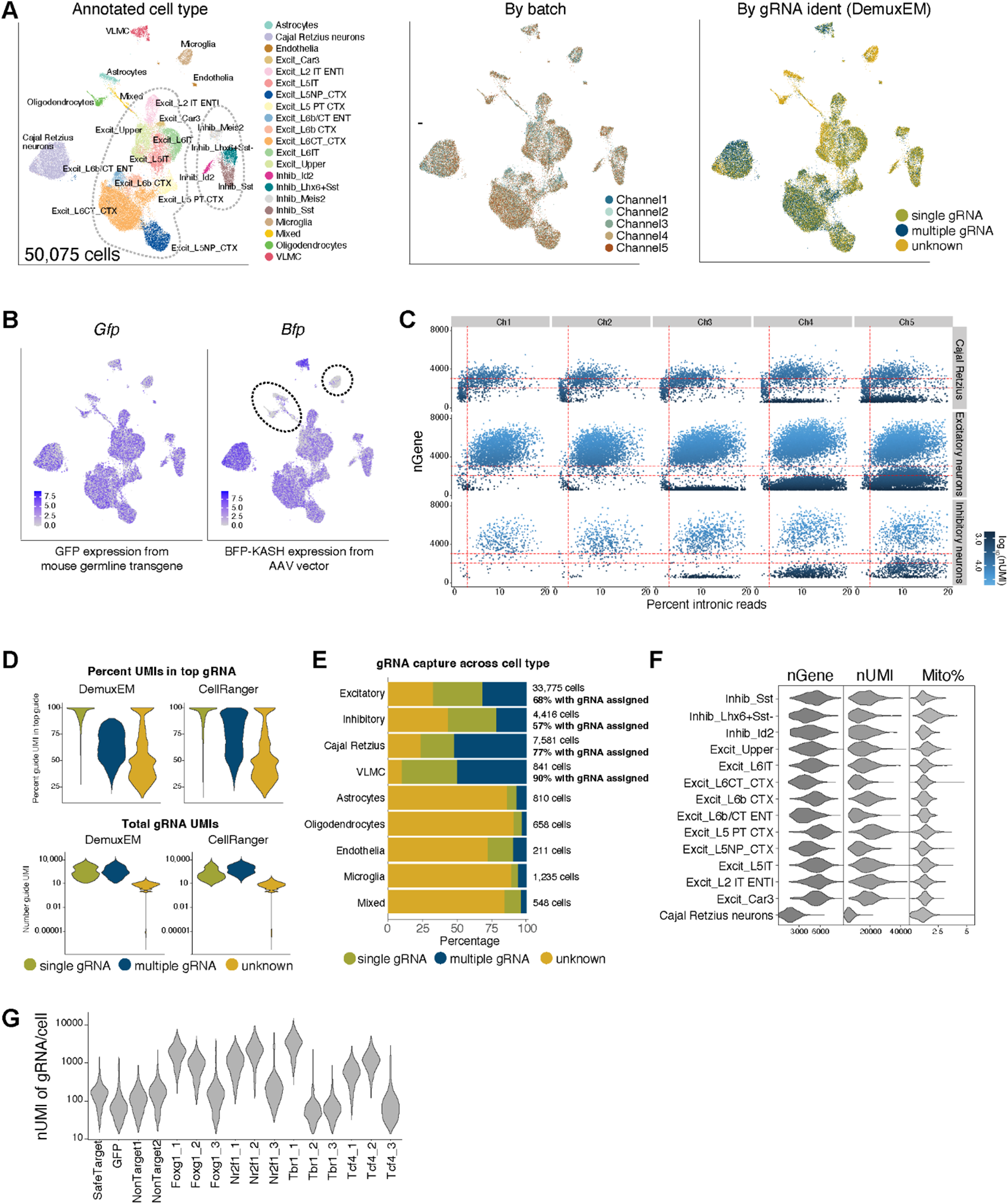
Quality control and gRNA identity annotation of the Perturb-seq data. (A) UMAP plots of all cells before downstream filtering, with each cell colored by annotated cell type (left), batch/channel (middle), and gRNA identity estimated by DemuxEM after down-sampling (right). (B) Feature plots of the log base 2 CPM expression of GFP (from the mouse transgene, expected to be detected in every cell) and BFP (only present in AAV transgene) across cell clusters. Dotted lines highlight the clusters with low BFP expression which were mostly glia. (C) Scatter plot showing the relationship of number of genes and percent of intron reads for quality control, with each dot representing a cell identified by Cell Ranger, the Y axis representing the number of genes, the X axis the percent of intronic reads in that cell. Lower left and lower right populations could be cytoplasmic or nuclei debris. Horizontal dotted lines represent 2,000 genes and 300 genes, vertical dotted line represents 3% intronic reads. (D) Violin plots comparing performance of DemuxEM (without down sampling) and Cell Ranger for assigning cells to gRNAs by evaluating percent of gRNA expression in top gRNA (top) or number of gRNA UMIs detected (bottom), across the annotated groups of cells with single, multiple, or no/unknown gRNA. (E) Bar plot of the proportion of cells assigned to one, no, or multiple gRNAs by down-sampling and DemuxEM. The measured gRNA capture efficiencies vary across major cell types. gRNAs were identified at a high rate in neurons while lower in non-neuron cells. (F) Violin plots showing the number of expressed genes per cell, number of UMIs per cell, and percentage of mitochondrial reads per cell in different cortical population. (G) Violin plot showing large differences in the number of gRNA UMIs recovered across perturbation groups.

**Figure S6.**
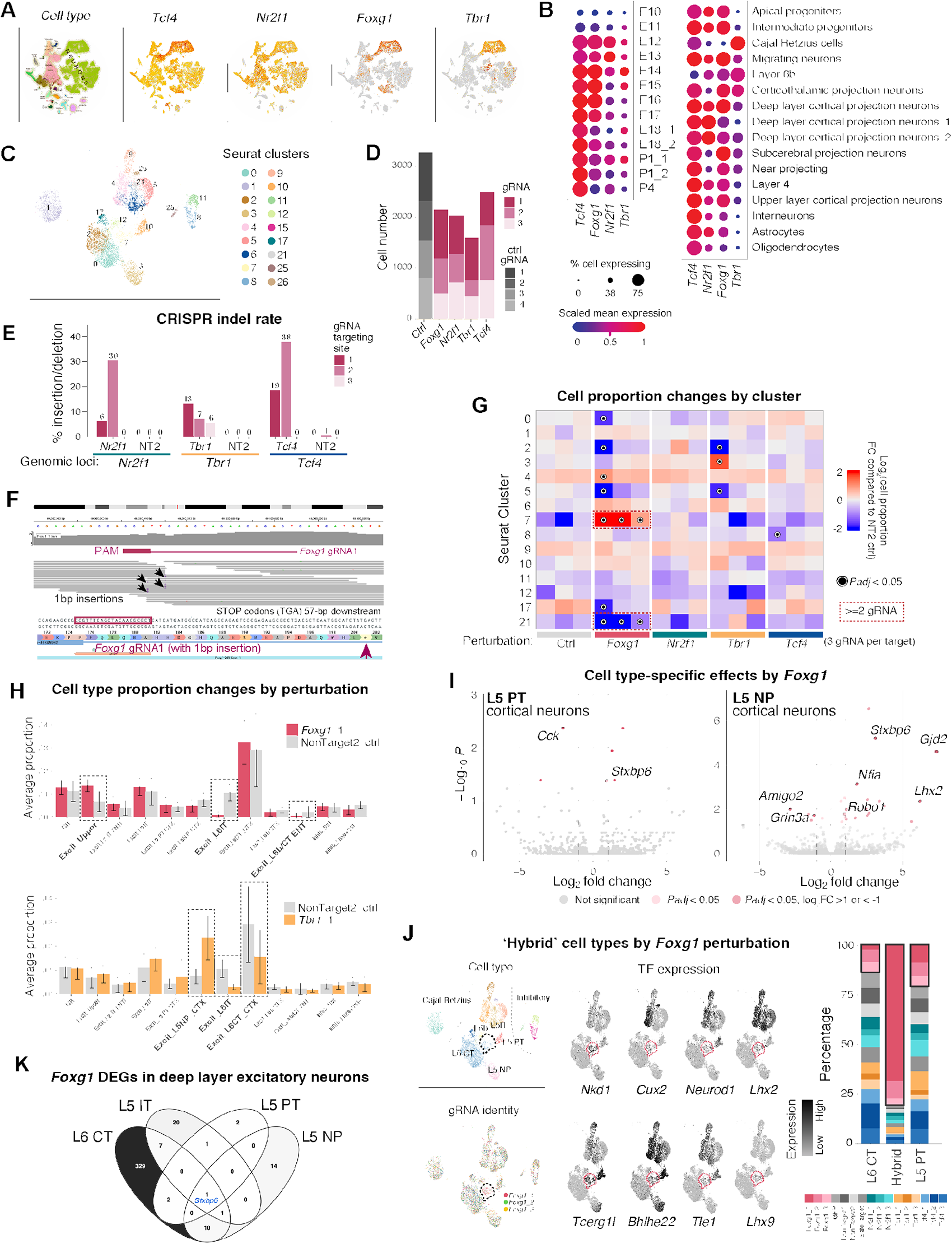
*In vivo* Perturb-seq identified cell type-specific changes across perturbations of transcription factors in the developing brain. (A) UMAP of the expression pattern of the 4 transcription factors (TFs) across cell types in mouse embryonic and perinatal development, colored by gene expression from La Manno et al. (generated from mousebrain.org). Two TFs are specifically expressed in certain cell types, and two TFs are expressed broadly. (B) Dot plot of the gene expression of the 4 TFs across cell types and time windows from Di Bella et al. Dot size corresponds to % of cells expressing that gene, color to normalized expression. (C) UMAP plot of filtered cells with a single perturbation, with each cell colored by Seurat cluster. (D) Bar plot of the cell number of each perturbation group containing one gRNA. The X axis is the gene targeted by a given guide, the color represents the guide number. (E) Bar plot of the percentage of cells with insertion/deletion (indel) in *Nr2f1, Tbr1* and *Tcf4* gRNA targeting area by perturbation or control NT2, extracted from scRNA-seq data. Gene-targeting gRNAs had variable efficiency while control gRNAs did not induce insertion/deletion on these loci. (F) IGV view of example reads showing *Foxg1* gRNA1 perturbation effects in inducing premature STOP codons 57 bp downstream, detected from the scRNA-seq data. (G) Heatmap showing cell cluster proportion changes by each perturbation, compared to controls. Color represents the log of cell proportion fold change compared to NT2 controls; black rings highlight FDR adjusted *P*-value<0.05, and red boxes highlight changes supported by at least 2 gRNAs. (H) Bar plots of the cell type proportion in *Foxg1* and *Tbr1* perturbations compared to NT2 controls. Error bars represent 95% confidence intervals, cell types with significant proportion change (*P*-adj<0.05) are in bold and labeled with black dotted line boxes. (I) Volcano plots of cell type-specific differential gene expression upon *Foxg1* gRNA1 perturbation in L5 PT and L5 NP neurons. (J) Perturbation of *Foxg1* led to a hybrid neuronal cell type. Left: UMAP plots of cell type and gRNA identity; dotted line highlights the hybrid subcluster (cluster 7). Middle: zoomed UMAP plots showing expression levels of mis-expressed transcription factors in the hybrid subcluster (pink dotted line). Right: proportion of perturbation groups in each of the three cell clusters. Black boxes highlight the three *Foxg1* perturbation groups. (K) Venn diagram of the number of overlapped differentially expressed genes (DEGs) by *Foxg1* gRNA1 perturbation across L6CT, L5 IT, L5 PT and L5 NP, with *Stxbp6* being the shared DEG in all 4 cell types.

