## Supplementary figures and images for "Massively parallel *in vivo* Perturb-seq reveals cell type-specific transcriptional networks in cortical development"

### Fig. S1

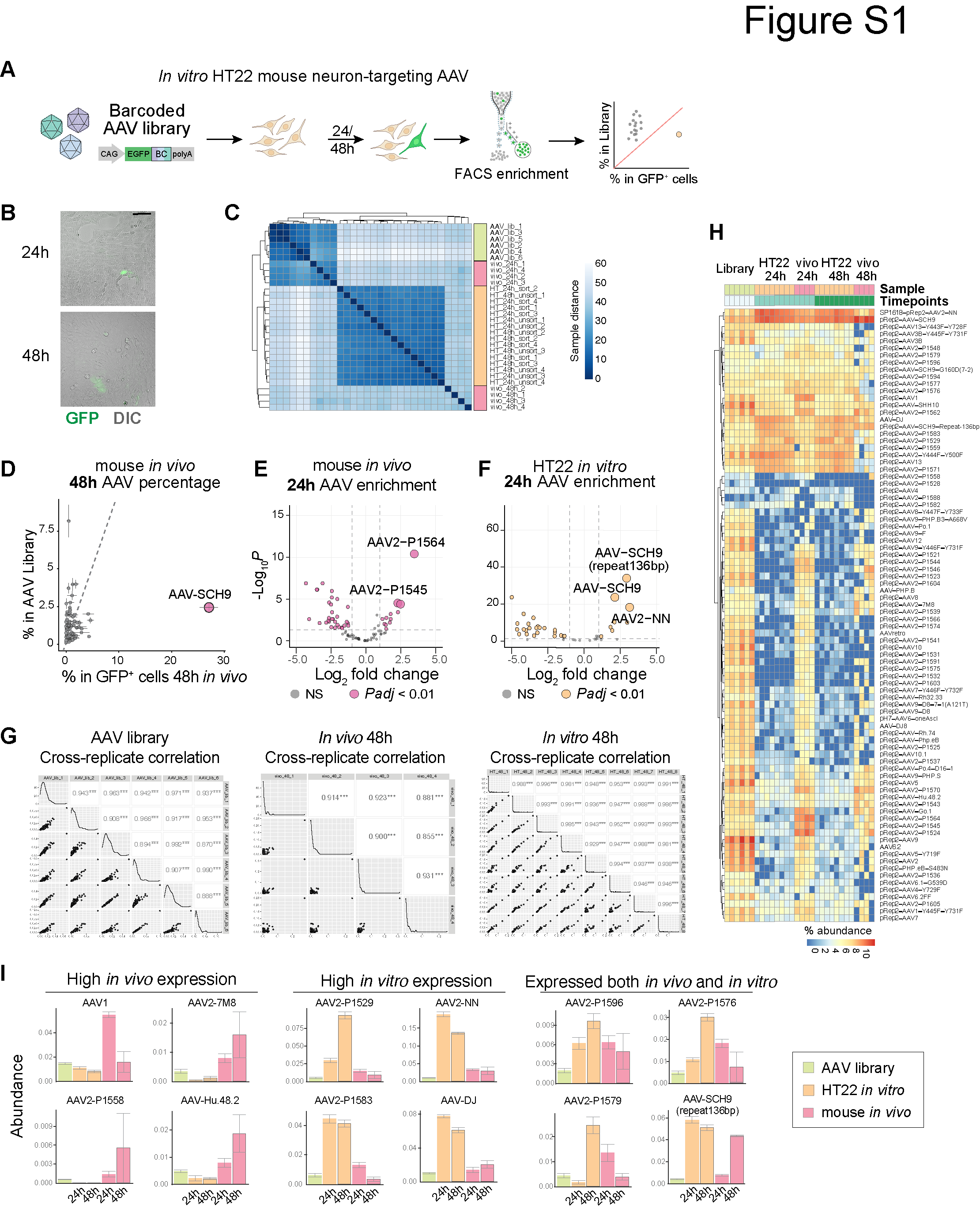

### Fig. S2

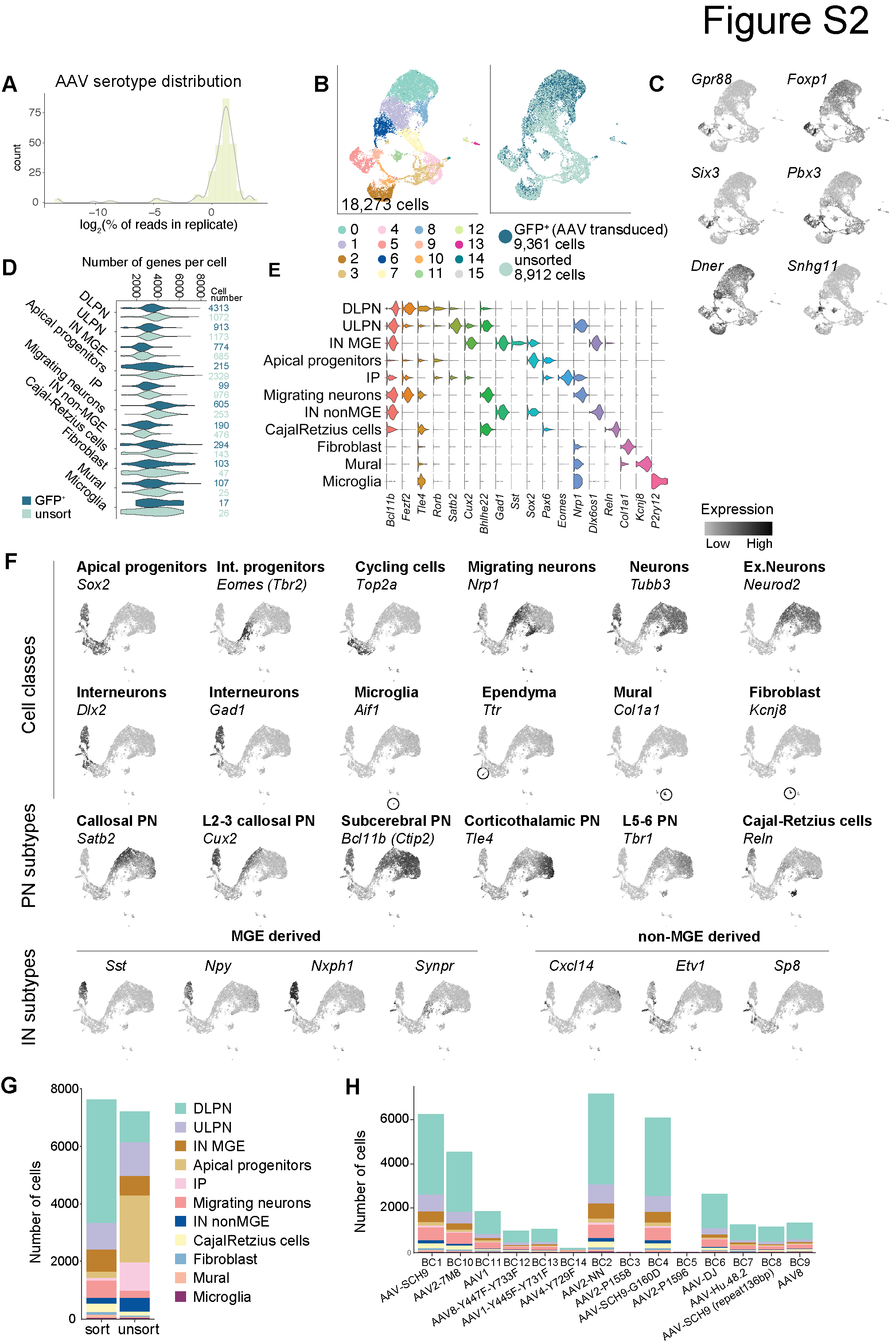

### Fig. S3

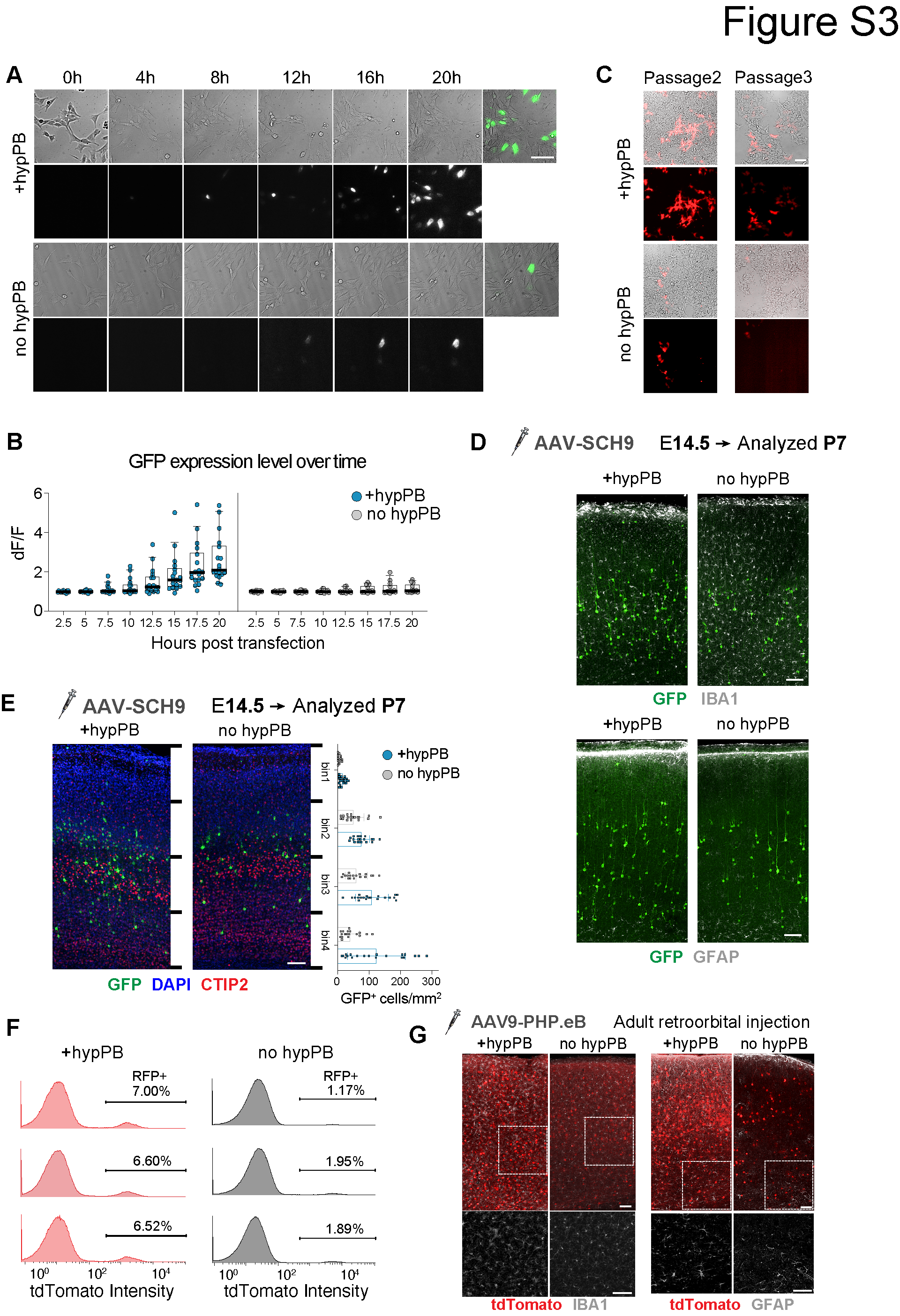

### Fig. S4

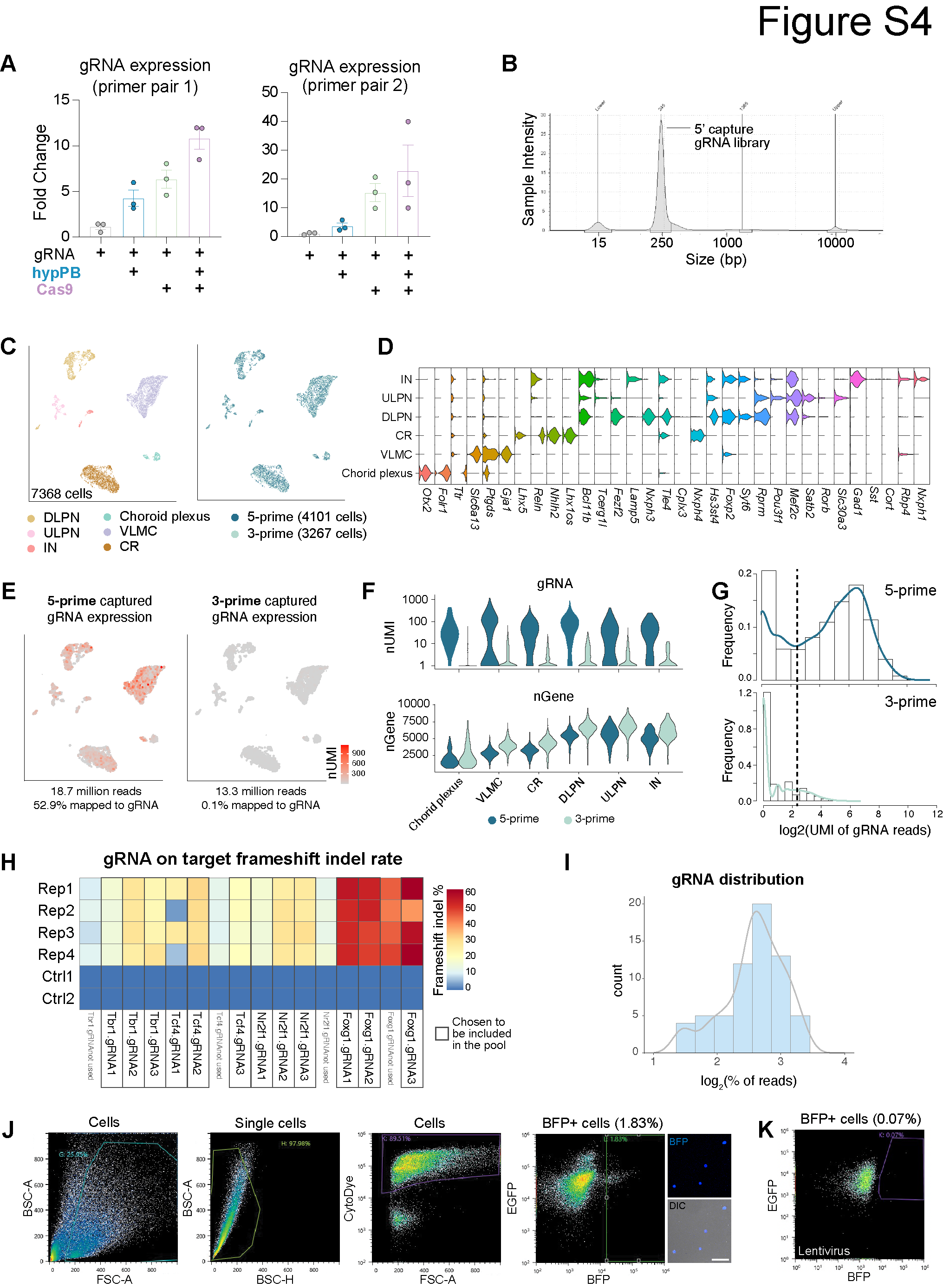

### Fig. S5

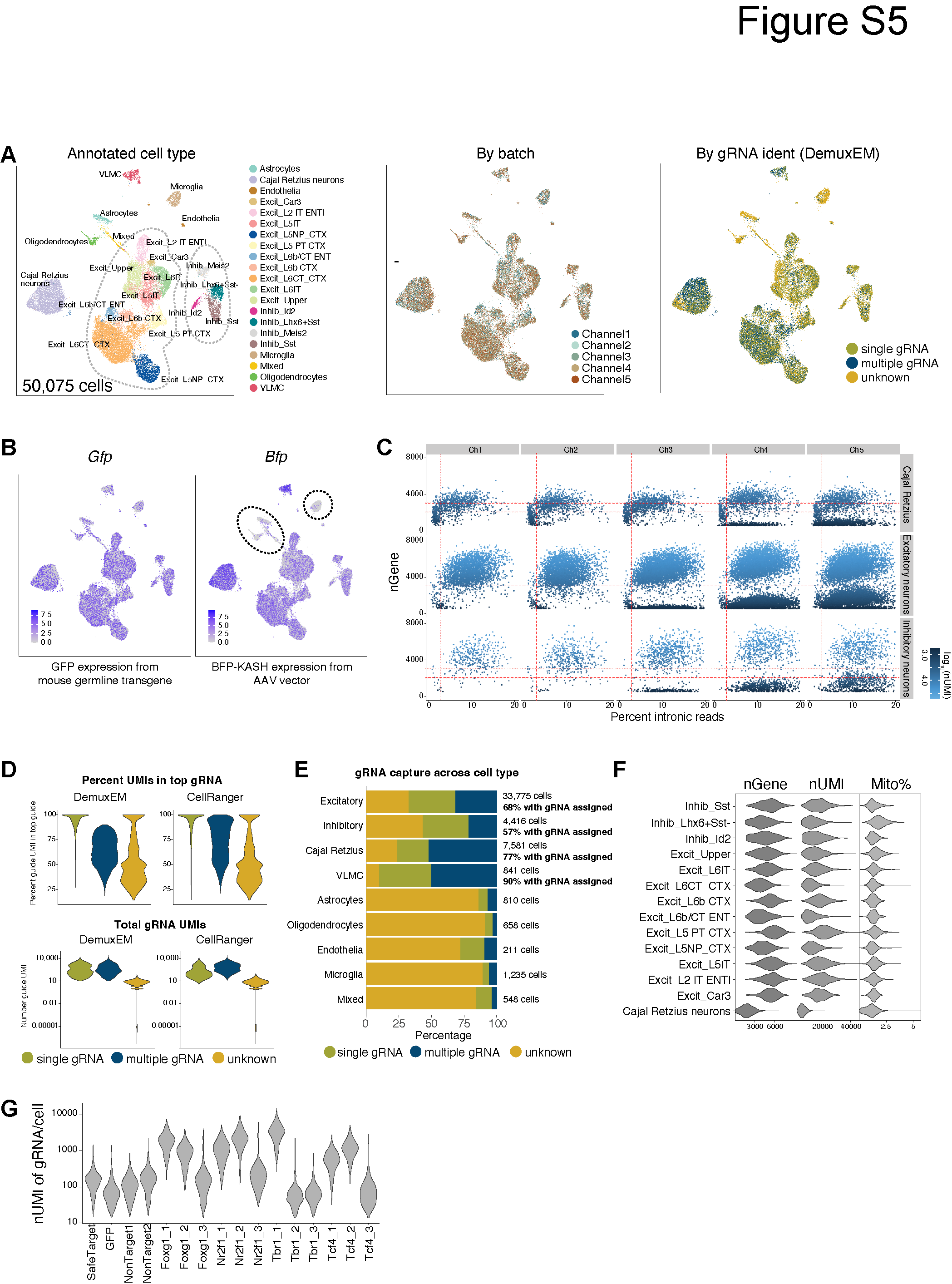

### Fig. S6

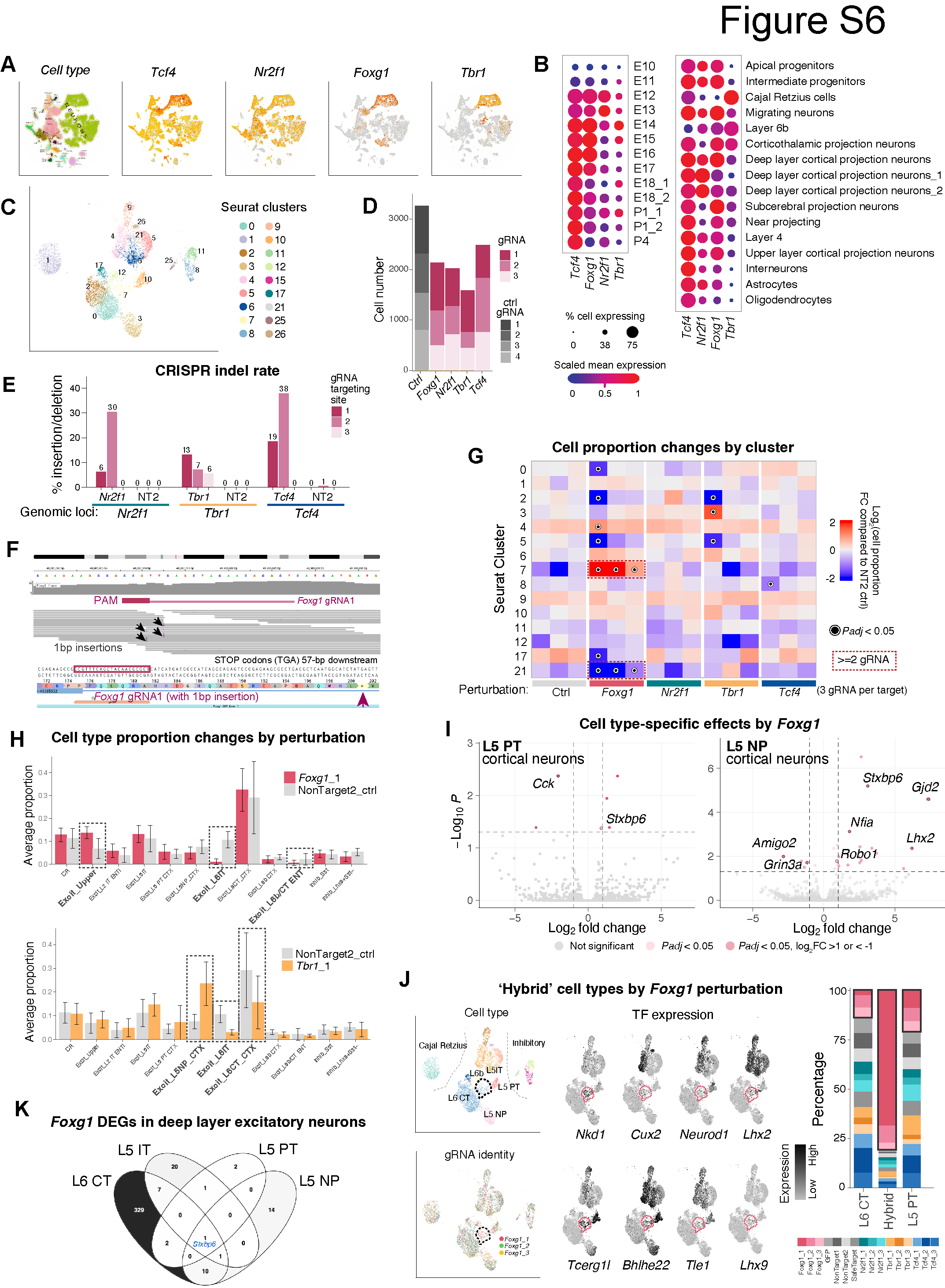
